# Subcellular orchestration of alpha-synuclein variants in Parkinson’s disease brains revealed by 3D multicolor STED microscopy

**DOI:** 10.1101/470476

**Authors:** Tim E. Moors, Christina A. Maat, Daniel Niedieker, Daniel Mona, Dennis Petersen, Evelien Timmermans-Huisman, Jeroen Kole, Samir F. El-Mashtoly, Liz Spycher, Wagner Zago, Robin Barbour, Olaf Mundigl, Klaus Kaluza, Sylwia Huber, Melanie N. Hug, Thomas Kremer, Mirko Ritter, Sebastian Dziadek, Jeroen J.G. Geurts, Klaus Gerwert, Markus Britschgi, Wilma D.J. van de Berg

**Author notes:** Corresponding authors: Wilma D.J. van de Berg, PhD, Dept. of Anatomy & Neurosciences, chair section Clinical Neuroanatomy and Biobanking, Amsterdam Neuroscience, Amsterdam UMC, location VU University Medical Center O2 building, room 13 E11, De Boelelaan 1108, 1081 HZ Amsterdam, the Netherlands.nr; Markus Britschgi, PhD, Roche Pharma Research and Early Development; Neuroscience, Ophthalmology, and Rare Diseases Discovery and Translational Area, Roche Innovation Center Basel, Grenzacherstrasse 124, CH – 4070, Basel, Switzerland. nr.

## Abstract

Post-translational modifications of alpha-synuclein (aSyn), particularly phosphorylation at Serine 129 (Ser129-p) and truncation of its C-terminus (CTT), have been implicated in Parkinson’s disease (PD) pathology. To gain more insight in the relevance of Ser129-p and CTT aSyn under physiological and pathological conditions, we investigated their subcellular distribution patterns in normal aged and PD brains using highly-selective antibodies in combination with 3D multicolor STED microscopy. We show that CTT aSyn localizes in mitochondria in PD patients and controls, whereas the organization of Ser129-p in a cytoplasmic network is strongly associated with pathology. Nigral Lewy bodies show an onion skin-like architecture, with a structured framework of Ser129-p aSyn and neurofilaments encapsulating CTT aSyn in their core, which displayed high content of proteins and lipids by label-free CARS microscopy. The subcellular phenotypes of antibody-labeled pathology identified in this study provide evidence for a crucial role of Ser129-p aSyn in Lewy body formation.

## Introduction

The presence of cellular inclusions – termed Lewy Bodies (LBs) and Lewy Neurites (LNs) – in predilected brain regions pathologically defines Parkinson’s disease (PD) and dementia with Lewy bodies (DLB). LBs and LNs are defined as eosinophilic inclusions with different morphologies, typically dependent on brain region (brainstem, limbic or cortical) ^1, 2^. The mechanism determining the formation and morphology of these inclusions remains elusive. LBs and LNs are strongly immunoreactive for alpha-synuclein (aSyn), which is one of their major components ^3^. aSyn is a 14kDa protein ubiquitously and highly expressed in neurons under physiological conditions. Its enrichment in presynaptic terminals has been established ^4–7^, while more recent studies have reported additional intraneuronal localizations for aSyn, including mitochondria, endoplasmatic reticulum (ER) and Golgi ^7^. The primary sequence of aSyn contains 140 amino acids, and is composed of three distinct domains. An important role has been proposed for the lipophilic N-terminus (NT) and non-amyloid-β component domain (NAC domain) in the interaction of aSyn with lipid membranes ^7, 8^, while the residues 96-140 encompass the negatively charged, acidic C-terminus (CT) of aSyn for which important roles have been proposed in the interaction of aSyn with other proteins or metal ions ^9^. The CT further harbors the majority of sites where aSyn can be post-translationally modified (PTM) ^10^.

The list of aSyn PTMs detected in the human brain has grown extensively in recent years, which highlights the physicochemical and structural flexibility of aSyn ^11, 12^. Some of these PTMs have been implicated in PD pathology - in particular the phosphorylation at Serine 129 (Ser129-p) and truncations of the C-terminus (CTT). Ser129-p aSyn and different CTT fragments of aSyn were identified in pathology-associated fractions of the DLB brain using mass spectrometry and immuno-based biochemical assays ^13–15^. Among the CTT variants most consistently identified in human brain tissue are the truncations at Asp-119 and Asn-122 ^14–16^. Although Ser129-p and CTTaSyn can be detected in small amounts under physiological circumstances^17^, these PTMs are enriched under pathological conditions ^13, 14, 16, 18, 19^. Immunohistochemical analyses in postmortem brain tissue of DLB patients and also aSyn transgenic mouse brains have pointed to a potential role of 122CTT in axonal and synaptic degeneration, which was ameliorated by blocking of calpain-mediated cleavage of aSyn by overexpressing calpastatin in aSyn transgenic mice ^20^. A laminar appearance of LBs with tyrosine hydroxylase (TH) and ubiquitin in the inner core, surrounded by 122CTT aSyn and Ser129-p aSyn in inner and outer layers, respectively, was previously described in the substantia nigra (SN) of subjects with incidential Lewy body disease (iLBD) and PD patients^21^. There results suggest that these forms of aSyn play a central role in the formation of LBs. Experimental studies have further suggested an important role for CTT in aSyn aggregation, as enhanced fibril formation was reported for this PTM with recombinant aSyn *in vitro* ^16, 22–25^. However, the exact role of CT modification by either phosphorylation or truncation in aSyn aggregation and toxicity, remains subject of active debate ^26^.

A great interest has emerged for PTM aSyn as a potential biomarker and therapeutic target for PD ^27, 28^, and led to the development of new research tools, including antibodies specifically directed against Ser129-p and CTT aSyn. Although such antibodies were reported to show immunoreactivity in LBs and LNs ^13, 14, 21^, only little is known about their detailed immunoreactivity patterns in the human brain. More information on the subcellular distribution of Ser129-p and CTT aSyn is crucial for a better understanding of their relevance in PD pathology, and therefore highly relevant for ongoing and upcoming immunovaccination therapies targeting different aSyn species. In this study, we aim to define the manifestation of Ser129-p and CTT aSyn in neurons under physiological and pathological conditions. For this purpose, we mapped subcellular immunoreactivity patterns of highly selective antibodies directed against Ser129-p and CTT aSyn in postmortem brain tissue of clinically diagnosed and neuropathologically verified PD patients, as well as donors with iLBD and aged non-neurological subjects, using high-resolution 3D confocal scanning laser microscopy (CSLM) and stimulated emission depletion (STED) microscopy.

Our results provide detailed insights into antibody-labeled subcellular pathology in PD, demonstrating a systematic onion skin-like architecture of nigral LBs composed of layers enriched for specific aSyn epitopes, in line with previous results from Prasad et al^21^. In extension of this previous finding, we discovered that Ser129-p aSyn at the periphery of such onion-skin type LBs is embedded in a structured cage-like framework of cytoskeletal components such as intermediate neurofilaments. Results of label-free coherent anti-Stokes Raman scattering (CARS) microscopy confirm the presence of increased lipid and protein contents in the core of a-Syn immunopositive inclusions. Together, these observations suggest the encapsulation of aggregated proteins and lipids in the core of LBs. Cytoplasmic CTT reactivity is associated with mitochondria in PD patients as well as controls, while Ser129-p aSyn is organized in a cytoplasmic network in neurons specifically in brains with LB pathology. The presence of this cytoplasmic network in neurons without inclusions in patients with early PD stages and incidental LB disease (Braak 3) suggests that this feature is an early subcellular manifestation of aSyn pathology. Based on our observations in the different experiments in this study, we identified a subset of subcellular phenotypes associated with pathology, which possibly reflect different maturation stages of Lewy pathology. Our data suggest extensive cellular regulation of Lewy body formation and maturation, and support a key role for Ser129-p aSyn in this process.

## Results

### Antibodies against aSyn protein variants highlight the heterogeneous nature of Lewy pathology

In accordance with existing literature ^13, 14, 21^, immunohistochemical (IHC) stainings using antibodies (specified in Supplementary Tables 1 and 5, and Supplementary Figures 9 and 10) selectively raised against truncated aSyn species (119 and 122 CTT aSyn) and Ser129-p aSyn labeled a variety of pathology-associated morphologies in post-mortem brain tissue from clinically diagnosed and neuropathologically confirmed advanced PD patients (Braak 5/6; Supplementary Table 2), including LBs and LNs. These neuronal inclusions were also detected using antibodies directed against epitopes within specific domains (CT, NT and NAC domain), while this group of antibodies revealed synaptic-like staining profiles in addition, most prominently in the hippocampus and transentorhinal cortex. Representative images of neuronal aSyn-positive inclusions labeled with antibodies against different aSyn epitopes, taken in the substantia nigra (SN), hippocampus and transentorhinal cortex of PD patients, are shown in Supplementary Figure 1, together with KM-51, an antibody commonly used for neuropathological diagnosis ^29^.

Lewy pathology in PD has been described to display substantial morphological heterogeneity, is amongst others determined by size of the inclusions, brain region, and specific cell type ^1^. When focusing on perinuclear aSyn-immunopositive intracytoplasmic inclusion bodies in the different analyzed brain regions (SN, hippocampus, transentorhinal cortex), we defined two main types of morphologies based on their IHC labeling profiles for antibodies against aSyn. In a large subset of spherical LBs in the SN, immunoreactivity for aSyn antibodies revealed a ring-shaped appearance, i.e. with a strong immunopositive band surrounding a central – weakly or unstained - core (Supplementary Figure 1A). Inspection in adjacent brain sections of the same patients suggested that this ring-shaped appearance of LBs was most clearly visualized by antibodies against Ser129-p aSyn and CT aSyn (Supplementary Figure 1A, xiv, xxvi). Antibodies directed against other aSyn epitopes generally revealed less contrast between core and immunoreactive ring. For certain antibodies, an area of weaker immunoreactivity surrounding the strongest immunopositive portion of LBs could be observed (f.i. Supplementary Figure 1A, xviii). The ring-shaped, lamellar appearance of midbrain LBs has been described with antibodies against aSyn in literature^21^, and a subset of these morphologies were shown to represent the eosinophilic ‘classical LBs’ unambiguously identified by hematoxylin and eosin stainings^2^. Peripheral immunoreactivity surrounding a weakly stained central core could also be observed in a subset of the dystrophic LNs in the SN (Supplementary Figure 1B).

Other intracytoplasmic aSyn-immunopositive inclusions showed diffuse and uniform labeling throughout the structure. This IHC pattern was generally observed for limbic and cortical LBs in the hippocampal CA2 region and transentorhinal cortex, respectively, but also for certain inclusions in the SN ^1, 2^. In the latter region, a subset of inclusions with uniform aSyn labeling have previously been termed as pale-body like ^1, 2^. The shapes of inclusion bodies uniformly labeled for aSyn showed substantial heterogeneity, including globular and compact appearing structures (f.i. Supplementary Figure 1A, xxiii), as well as irregularly-shaped expansive-appearing cytoplasmic inclusions (f.i. Supplementary Figure 1A, i&iii), as has been previously described for cortical LBs ^1, 2, 30^.

### A subset of nigral LBs and LNs displays a consistent onion skin-like distribution of aSyn epitopes

To further assess the exact localization patterns of aSyn epitopes in different LB and LN morphologies, multiple immunofluorescent labelings were performed using antibodies selectively raised against different aSyn variants, in different combinations and stained features were inspected by high-resolution confocal and STED microscopy. Similar to our observations in brightfield microscopy, we observed a variety of aSyn-immunopositive features. As described before, a major distinction could be made between LB and LN morphologies that revealed either a ring-shaped appearance or a uniform distribution of Ser129-p aSyn immunoreactivity.

In the subset of ring-shaped LBs in the SN, we observed that Ser129-p aSyn, 119CTT and 122CTT only partially co-localized, using different combinations of antibodies (Supplementary Tables 1 and 5 and Supplementary Figures 9 and 10). In particular, Ser129-p immunopositive rings in nigral LBs localized consistently more towards their periphery as compared to staining patterns for 119CTT or 122CTT aSyn (Figure 1A, C, Supplementary Video 1,2). Interestingly, different localization patterns were also observed between antibodies against NAC, NT and CT residues 118-126 of aSyn, with CT aSyn immunoreactivity surrounding the other domains (Figure 1B, D, Supplementary Video 3).

**Fig 1:**
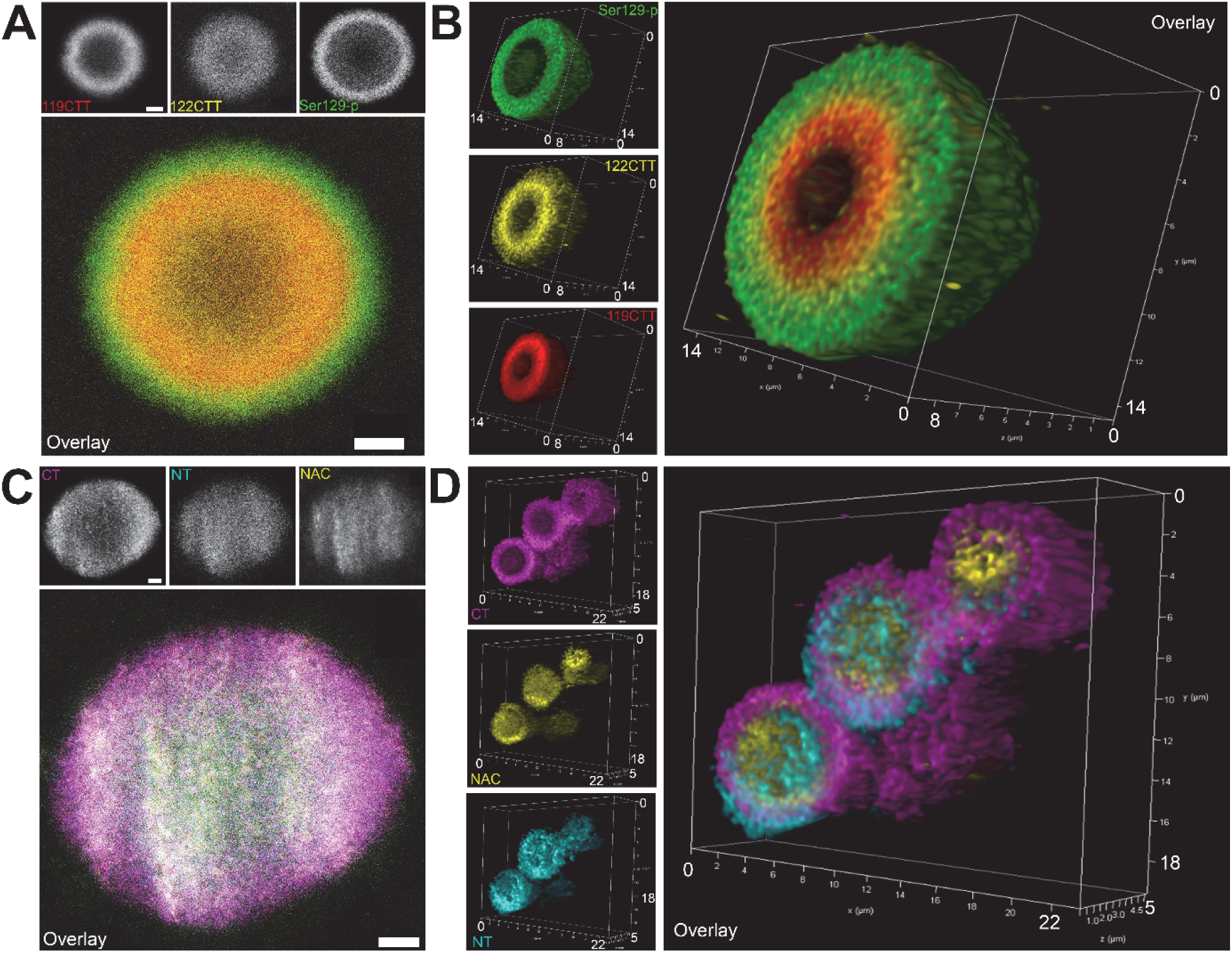
Lamellar distribution patterns of aSyn PTMs and aSyn domains in LBs. **A:** Triple labeling of aSyn PTMs: representative raw STED image of a nigral LBin patient PD7, showing immunoreactivity for Ser129-p aSyn at the periphery of the LB while 119CTT and 122 CTT aSyn are localized in the core of the structure. **B:** 3D reconstruction based on deconvolved CSLM images showing the lamellar distribution of different aSyn PTMs in a nigral LB in patient PD6. **C:** Triple labeling of different aSyn domains: raw STED image of a nigral LB, showing immunoreactivity for CT aSyn at the periphery of the LB and NT and NAC aSyn staining in the core of the structure, taken in the SN of patient PD8. **C:** 3D reconstruction based on deconvolved CSLM images showing different localization for different aSyn domains in a complex of three LBs in the SN of patient PD8. Scale bar in **A** and **C:** = 2 µm

When combining all these antibodies in one multiple labeling protocol, a gradual and distinct distribution of immunoreactivities in nigral LBs was observed (Figure 2A). The lamellar organization of different concentric rings together demonstrated an onion skin-like morphology (Figure 2E, Supplementary Video 4), with pronounced reactivity for CT and Ser129-p aSyn at the periphery, while antibodies against CTT aSyn, NT and NAC aSyn localized more towards the core of these structures (Figure 2A-C). These distribution differences were confirmed using a set of different antibodies directed against similar epitopes (Supplementary Figure 2; Supplementary Table 1). Converging DAPI reactivity was consistently observed at the core of midbrain-type LBs, although this signal was generally weaker than its staining intensity in neighboring cell nuclei (Figure 2B). 3D CSLM analyses showed that the lamellar build-up of this subset of LBs was present throughout the entire structure (Figure 2E, Supplementary Video 4). Interestingly, dystrophic LNs in the SN were frequently observed to contain compositions similar to these LBs (Figure 2D).

**Fig 2:**
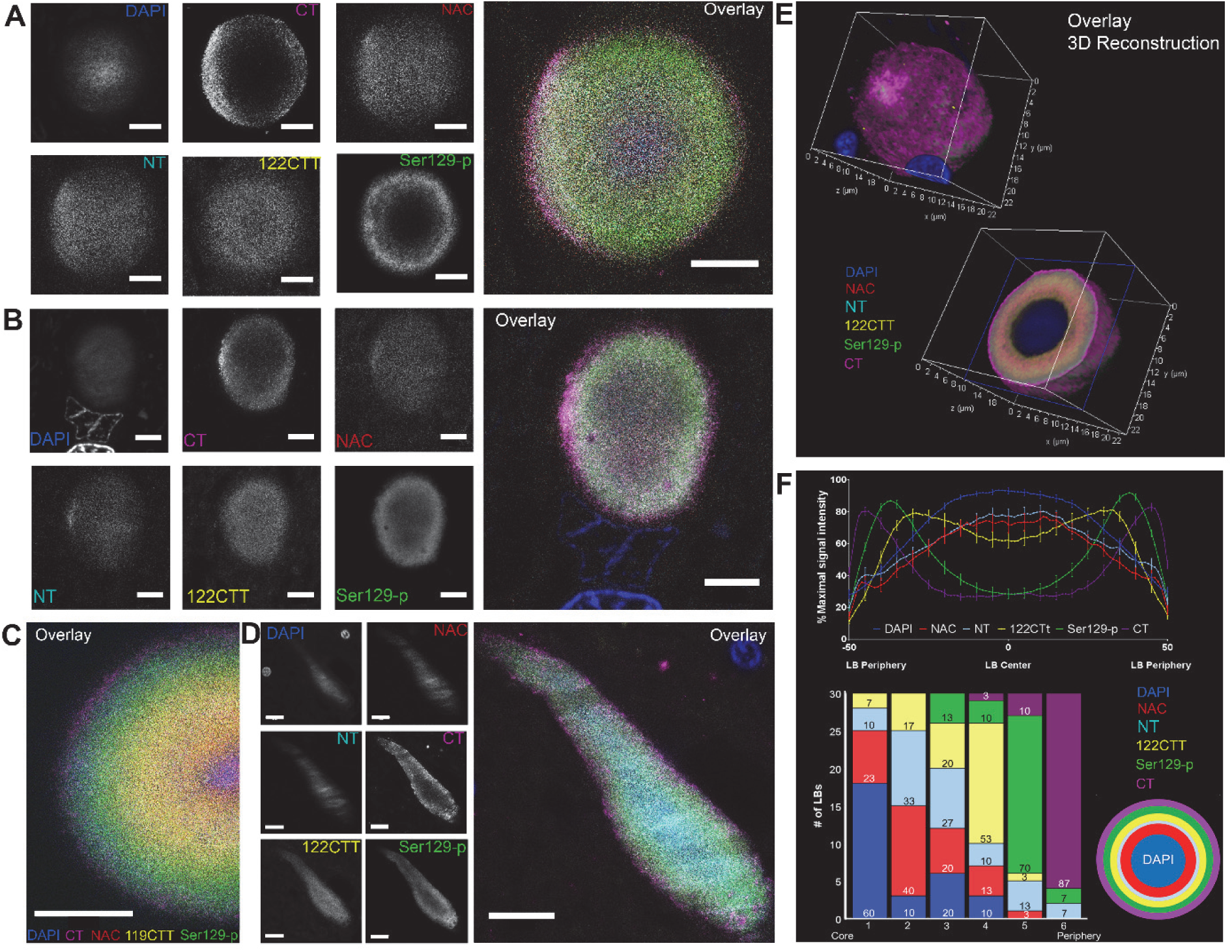
Onion skin-like orchestration of different aSyn epitopes in nigral LBs and LNs. **A, B:** Raw STED-images showing immunoreactivities for different aSyn epitopes in onion skin-type LBs in the SN of patient PD1 (**A)** and PD5 (**B**). Immunoreactivities for CT and Ser129-p aSyn are localized at the periphery of the structures, while NT, NAC and 122CTT reactivities were present mainly in their core. **C:** Raw STED image of protocol including an antibody against 119CTT aSyn, taken in the SN of patient PD5. **D:** Raw STED image of a dystrophic LN in the SN of patient PD7. **E:** 3D reconstruction of an entire nigral LB based on deconvolved CSLM images in patient PD1. **F:** Top: Average line profile (±SEM) for 30 onion skin-like LBs measured in the SN of 8 PD patients, showing a separation of peak intensities. Bottom: summary of rankings of peak intensity locations from core to periphery for the studied aSyn epitopes, highlighting peripheral localization of CT and Ser129-p aSyn. Right: a schematic depiction of the lamellar architecture of LBs as revealed by antibodies against different aSyn epitopes. Source data for this Figure are provided as a Source Data file. **A-C:** Scale bar = 5 µm; **D:** Scale bar = 10 µm.

To test the consistency of this observation, we semi-quantitatively examined 30 LBs in formalin-fixed paraffin embedded SN sections from 7 PD patients (Supplementary Table 3). LBs included for analysis contained a Ser129-p aSyn immunopositive ring-shaped appearance, were localized in the cytoplasm of neuromelanin-containing dopaminergic neurons, and had a diameter larger than 5µm (further explained in Methods). For each of the scanned LBs, relative signal intensities (as % of peak intensity within the same structure) were plotted per channel over a normalized LB diameter, in order to generate line profiles that visualize the distributions of aSyn epitopes in each LB. The average line profile over the 30 LBs showed a clear separation of peak intensity localizations for the different aSyn epitopes, revealing that this gradual difference in distributions was consistent in the LBs selected for analysis (Fig. 2F, upper panel). In addition, we determined the position in each LB where the peak intensity for each aSyn epitope was localized relative to the LB origin, which we ranked among the different epitopes (Fig. 2F, lower panel). A different distribution of peak intensities was observed and confirmed by statistical analysis of this data (χ^2^: 73.912 (4); p <0.0001; Figure 2F; Supplementary Table 3; post-hoc tests presented in Supplementary Table 4).

The peak intensities of 122CTT aSyn were localized at a more central position compared to Ser129-p aSyn in almost all (97%; Table S3) analyzed LBs. Moreover, immunoreactivity of CTT aSyn was localized more to the core of LBs than antibodies against res 118-126 of aSyn’s CT in 97% of the analyzed LBs (Table S3), which was even more localized to the periphery than Ser129-p aSyn (Figure 2F). This finding confirms that the antibodies against 122CTT and CT aSyn in our study detect different aSyn species, and indicate that most of aSyn with an intact 118-126 portion of the epitope aSyn is present at the extreme periphery of LBs, while a substantial portion of aSyn in the LB core contains shortened CT. This effect could be specific for truncations at the CT, as peak intensities for NAC and NT aSyn were found more towards the central portion of the LB (Fig. 2F). No differences were observed between LBs measured in different patients for our analysis.

Distribution patterns for different aSyn epitopes were also analyzed in cytoplasmic neuronal inclusions with uniform Ser129-p immunoreactivity without ring-shaped appearance, in the same sections (in case of SN) and in other sections of the same patients (hippocampus/transentorhinal cortex). A detailed look on these inclusion bodies showed a homogeneous labeling for Ser129-p aSyn, except for typical ‘cavities’ lacking Ser129-p reactivity (Supplementary Figure 3A,D). Although such inclusions were unambiguously stained for Ser129-p aSyn, antibodies against other aSyn epitopes (particularly 122CTT, NAC and NT aSyn) generally revealed weak signal intensities that were distributed diffusely throughout the inclusion, while immunoreactivity for DAPI was barely increased compared to the surrounding. As a result of this, these inclusions appeared relatively unstructured – e.g. no systematic distribution patterns for aSyn epitopes were observed(f.i. Supplementary Figure 3A, Supplementary Video 5). Taken together, our results show morphology-dependent distribution patterns for aSyn epitopes in Ser129-p aSyn-positive cytoplasmic and neuritic inclusions, and provide evidence for a consistent lamellar and onion skin-like 3D organization of a subset of midbrain LBs and LNs.

### Onion skin-type LBs contain a cytoskeletal framework associated with Ser129-p aSyn

The consistent lamellar architecture of midbrain onion skin-type LBs may suggest that this LB morphology is the result of extensive cellular organization. Major constituents in organizing cellular organelles and substructures are cytoskeletal proteins, which have been identified in LBs by immunohistochemical approaches before ^31^. In order to obtain more insight into their detailed distribution patterns in onion skin-type LBs and their association with aSyn immunoreactivity patterns, we applied additional multiple labeling experiments including markers for intermediate neurofilaments and beta-tubulin which were analyzed using 3D gated STED microscopy. Markers for intermediate neurofilament and beta-tubulin showed immunoreactivity mainly at the periphery of nigral onion-skin type LBs, where they were associated with the immunoreactive band for Ser129-p aSyn (Figure 3). Together with Ser129-p aSyn, cytoskeletal markers visualized a ‘cage-like’ framework formed by cytoskeletal components and Ser129-p aSyn at the peripheral portion of LBs (Figure 3D).

**Fig 3:**
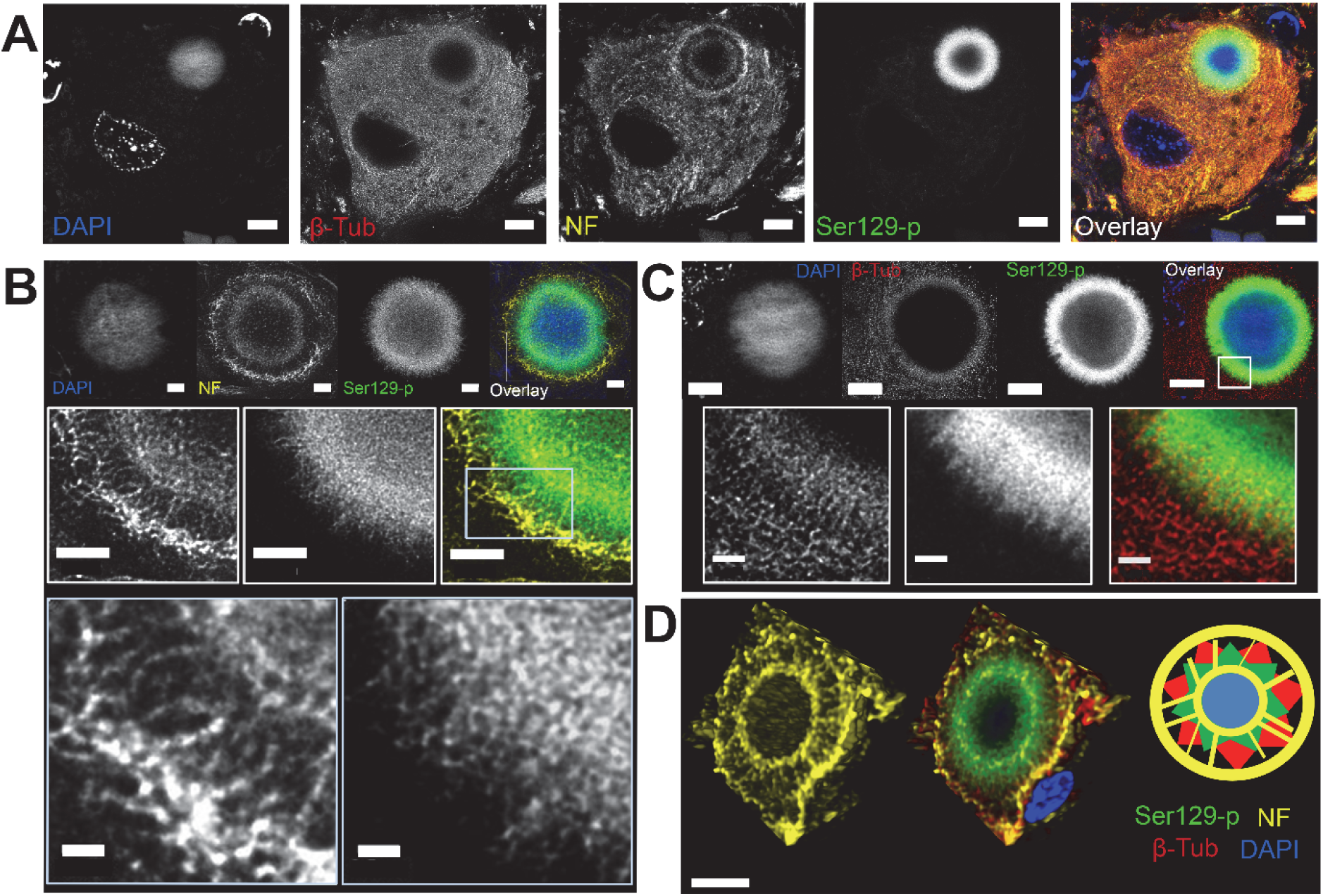
Ser129-p aSyn forms a cage-like framework with cytoskeletal components at the periphery of nigral LBs. **A:** Deconvolved STED image of neuromelanin-containing dopaminergic neuron in the SN with LB. Immunoreactivity for beta-tubulin and neurofilament (in two rings) is observed at the periphery of the LB. **B:** Deconvolved STED images showing the detailed structure of neurofilament in a onion skin-type LB at different magnifications. **C:** Deconvolved STED images showing detailed beta-tubulin reactivity at the periphery of a LB. **D:** Left: 3D reconstruction of the localization of Ser129-p aSyn and cytoskeletal components in a nigral LB, highlighting the wheel-like structure of neurofilament. Right: schematic summary of the results. NF: neurofilament; β-Tub: beta-tubulin. **A:** Scale bar = 5 µm; **B:** upper and middle row: scale bar = 2 µm; lower row: scale bar = 0.5 µm; **C:** Upper row: Scale bar = 5 µm, lower row: scale bar = 1 µm. **D:** Scale bar = 5 µm.

Although LBs are defined in brightfield microscopy as spherical smooth-edged inclusions, detailed STED imaging revealed that the outline of many LBs revealed irregular and radiating Ser129-p aSyn immunoreactivity patterns (Supplementary Figure 4). Beta-tubulin immunoreactivity showed substantial co-localization with Ser129-p aSyn at the LB periphery, and visualized similar radiating features, although even localized slightly more towards the outer LB portion (Figure 3B). Antibodies against intermediate neurofilaments, in contrast, revealed a remarkably structured organization at the periphery of LBs. In particular, two immunopositive rings were labeled in LBs: one ring localized at the central portion of the Ser129-p aSyn immunopositive band, while another ring surrounded the Ser129-p/beta tubulin signals. These rings are connected by neurofilament-immunoreactive elements giving rise to a structure resembling a wheel (Figure 3B). The detailed distribution of cytoskeletal components around the Ser129-p immunopositive band was also observed in 3D (Figure 3D, Supplementary Video 6-S8).

The wheel-like structure of neurofilaments at the peripheral portion of LBs was observed in many onion skin-type morphologies in the SN of all PD patients analyzed in this study, suggesting that this is a general feature of this LB-type (Supplementary Figure 5). For cytoplasmic inclusions uniformly labeled for Ser129-p aSyn and (dystrophic) LNs, the presence of cytoskeletal markers was less prominent, although (diffuse) immunoreactivity was occasionally observed in these morphologies. Together, this demonstrated that he organization of Ser129-p aSyn and cytoskeletal markers was characteristic for a specific subset of cytoplasmic inclusions, and may indicate that their recruitment and organization is an event related to certain maturation stages of LBs.

### Lipids and proteins are centralized in nigral aSyn-positive inclusion bodies

Together, our high-resolution microscopic observations revealed that in the heterogeneous landscape of aSyn pathology, a subset of morphologies reveals a consistent and structured ‘onion skin-like’ organization, pointing to the possibility that this LB morphology reflects an organized encapsulation of material in the LB core. Recently, we demonstrated that LBs contain increased protein and lipid levels using a label-free nonlinear optical imaging technique, CARS, on native brain tissue sections in combination with confocal microscopy ^32^. Here, we applied the a similar pipeline (Supplementary Figure 11) on sections of fresh-frozen midbrain tissue - including the SNpc - of 5 PD patients (Supplementary Table 2), to explore the distribution of lipid and protein contents in Ser129-p aSyn-immunopositive inclusions.

We included a total of 57 neuronal inclusion bodies (3 to 7 inclusion bodies per patient) immunopositive for Ser129-p aSyn with a diameter larger than 5µm in the SNpc of these sections. In line with our observations in brightfield microscopy, CARS data of these inclusions revealed substantial heterogeneity in the chemical composition of aSyn inclusions, both within and between patients (Figure 4E). Among the scanned inclusions were structures displaying higher levels for both proteins and lipids, inclusions with higher protein levels without increased lipid levels, and inclusions without detectable differences in protein and lipid levels compared to the environment (Figure 4A-C). The majority of scanned sctructures showed higher protein levels (37 out of 57) as compared to the direct environment, while higher lipid content was detected by CARS in 20 out of 57 inclusions (Figure 4E).

**Fig 4:**
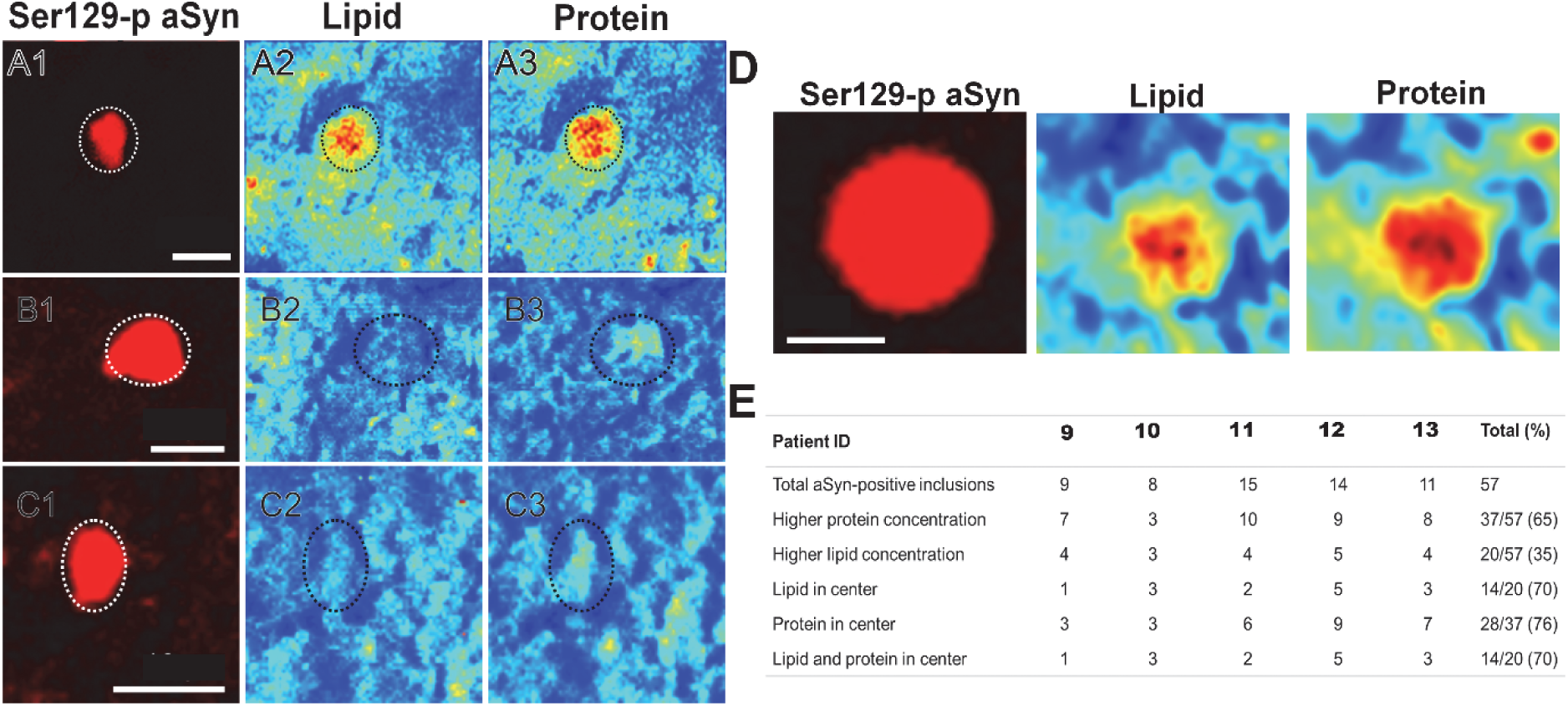
Protein and lipid distribution of nigral LBs. **A-C:** Different compositions of LBs as identified by CARS microscopy. The fluorescence images of Ser129-p aSyn of different LB types are depicted (first column). CARS signal intensities at 2850 cm^−1^ and 2930 cm^−1^ shows lipid (second column) and protein (third column) distributions, respectively. Low CARS-intensities are depicted in blue, whereas high intensities are depicted in red. LBs with different compositions were identified: LBs with high CARS intensities for proteins and lipids compared to the direct environment (A1-3), with high CARS intensity for proteins but not lipids (B1-3), and with low CARS intensity for proteins and lipids (C1-3). **D:** Representative image of a LB with high protein and lipid signal centralized in the structure. **E:** Numbers and proportions of nigral LBs with high (centralized) lipids or proteins per patient. In total 57 LBs were observed in 5 PDD patients (n=3-7 per patient) of which 37 showed high protein concentration and 20 showed high lipid concentration compared to surrounding tissue. In total 14 out of 20 with high lipid concentration showed lipids mainly in the center, whereas 28 out of 37 showed mainly proteins in the center.**A-C:** Scale bar = 10 µm; **D:** Scale bar = 5µm.

Despite this heterogeneity and the limited total number of inclusions scanned, some recurring patterns could be observed in the distribution of lipid and proteins in the LBs. When considering the subset of inclusions in which increased lipid content was detected, we observed that the increase in lipids was more pronounced in their central portion (in 14 out of 20 inclusions; Figure 4E). This pattern was also observed for proteins, as 28 out of 37 inclusions showed increased protein content mainly in the center. Interestingly, in all 14 inclusions that demonstrated centralized lipid content, this was accompanied by increased protein content in the core. Overall, our results show that in the subset of aSyn-positive inclusions that shows increased protein and lipid content compared to the surrounding, these components are predominantly in the central portion of this structure, supporting the idea that LBs represent organized encapsulations of accumulated cellular material.

### Distinct cytoplasmic manifestations of Ser129-p and CTT aSyn

Apart from their localization in aSyn-immunopositive inclusion bodies, Ser129-p and 122CTT aSyn antibodies also revealed cytoplasmic immunoreactivity in neurons outside of these structures. Immunoreactivity for 119CTT was specific for pathological inclusions with limited immunoreactivity in the cytoplasm. The appearance of cytoplasmic reactivity was different for Ser129-p and 122CTT aSyn (Figure 5A). In particular, 122CTT aSyn revealed many immunoreactive punctae throughout the neuronal cytoplasm, while Ser129-p aSyn immunoreactivity visualized an intracytoplasmic network surrounding a nucleus lacking immunoreactivity (Figure 5A). This cytoplasmic network was sometimes in continuation with similar Ser129-p immunoreactive features in the proximal portion of the neuronal processes. Limited co-localization was observed between the 122CTT immunopositive punctae and the cytoplasmic network immunopositive for Ser129-p aSyn.

**Fig 5:**
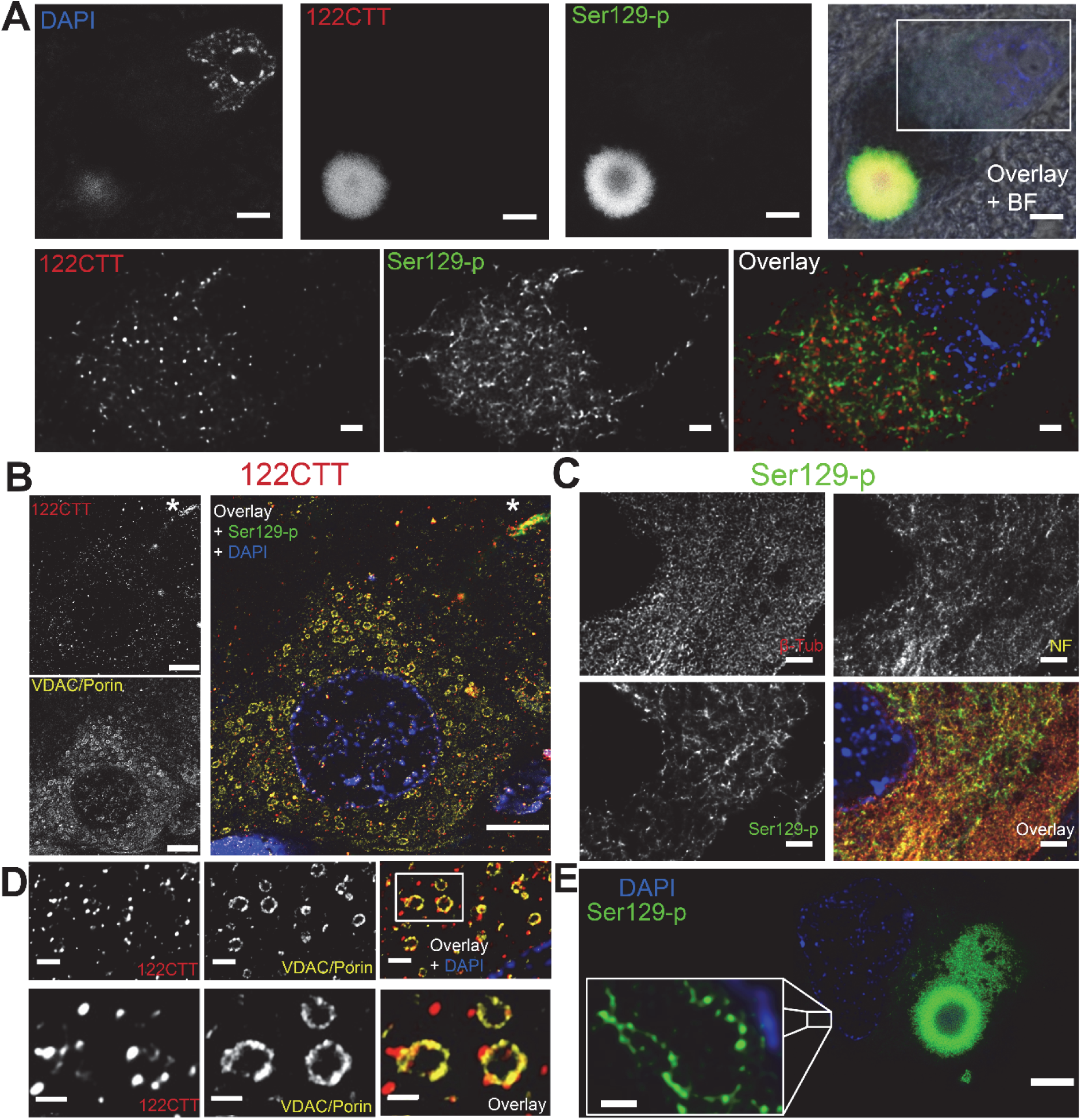
Subcellular manifestation of features containing 122CTT and Ser129-p aSyn. **A:** Overview of a neuromelanin-containing dopaminergic neuron in the SN with a LB (upper row) and a zoom-in on its cytoplasm (lower row). Signal intensity for 122CTT and Ser129-p aSyn was highest in the LB, suggesting an enrichment for these PTMs in pathological inclusions. However, cytoplasmic reactivity could also be observed, showing different manifestations for 122CTT and Ser129-p aSyn. **B:** Cytoplasmic 122CTT immunopositive punctae showed association with staining patterns for VDAC/Porin-immunopositive mitochondria in the hippocampal CA2 of a PD patient. Staining was also observed in a LN (indicated with an asterix). **C:** Ser129-p aSyn immunopositive network showed limited overlap with intracytoplasmic networks visualized by markers for beta-tubulin and neurofilament in the cytoplasm of a melanin-containing dopaminergic neuron in the SN of a PD patient. **D:** Localization of 122CTT immunopositive punctae at the outer membrane of mitochondria in the hippocampal CA2 of a PD patient at different magnifications. **E:** Dopaminergic neuron in the SN containing a combination of expansive-appearing inclusion and onion-skin type LB surrounded by a Ser129-p aSyn immunoreactive network. Insert: zoom in of Ser129-p aSyn immunoreactive elements with a diameter of ∼70nm. **A:** Deconvolved CSLM images; **B-E:** deconvolved STED images. **A:** Upper row: scale bar = 5 µm; lower row: scale bar = 2 µm; **B:** scale bar = 5 µm; **C:** scale bar = 2 µm; **D:** upper row: scale bar = 1 µm; lower row: scale bar = 0.5 µm; **E:** main figure: scale bar = 5 µm; inset: scale bar = 0.5 µm.

#### 122CTT aSyn punctae are associated with mitochondria

122CTT immunopositive punctae were not only observed in PD patients, but also in brain tissue sections of donors without Lewy pathology (Supplementary Figure 7A). The pattern appeared more pronounced in the hippocampus and transentorhinal cortex compared to the SN (Figure 5B). We observed the punctate reactivity pattern of 122CTT aSyn using different antibodies against this epitope (syn105 and asyn-134), while this pattern was not observed for 119CTT aSyn. These 122CTT aSyn-reactive punctae were more pronounced in the cytoplasm of neurons compared to the environment. Analysis with subcellular markers revealed that a subset of the 122CTT-reactive punctae in the cytoplasm of neurons co-localized with mitochondrial morphologies immunoreactive for Porin/VDAC reactivity, which is a marker for the outer membrane of mitochondria (Figure 5B). This association between CTT aSyn and mitochondria was also observed in 3D (Supplementary Video 10). Together, our findings suggest widespread 122CTT aSyn expression in the brain of donors with and without PD, particularly in the neuronal cytoplasm where 122CTT aSyn is possibly associated with mitochondria.

#### Ser129-p aSyn reveals disease-associated intracytoplasmic network

The Ser129-p aSyn immunopositive cytoplasmic network was frequently visible in large neuromelanin-containing neurons in the SN (Figure 5C), but also in other cell types, such as pyramidal neurons in the transentorhinal cortex. Most often, cytoplasmic Ser129-p aSyn immunoreactivity was observed in neurons containing an expansive-appearing inclusion uniformly stained for Ser129-p aSyn, and in a smaller fraction of neurons containing onion skin-type LBs. However, the cytoplasmic network was also observed in a subset of neuromelanin-containing neurons without apparent inclusion in PD patients (f.i. Figure 6A, 1). Other neuromelanin-containing neurons without inclusion in donors with PD did not reveal a Ser129-p aSyn immunopositive cytoplasmic network, while it was not observed in donors without Lewy pathology (Supplementary Figure 7B). The Ser129-p aSyn immunoreactive cytoplasmic network showed only limited overlap with other intracytoplasmic networks such as intermediate neurofilaments, beta-tubulin (Figure 5C) or endoplasmatic reticulum (ER; Supplementary Figure 6). This indicates that the alignment of Ser129-p positive elements is not simply explained by localization of Ser129-p aSyn to these networks. The observed diameter of the structures in the network was generally 70-80 nm and probably limited by the resolution of the applied scan settings. Importantly, the same feature was also identified in donors with iLBD or for PD patients staged as Braak 3, suggesting that the Ser129-p aSyn network manifests already at early stages of LB formation (Supplementary Figure 8, Supplementary Video 9) ^33^. Together, these observations suggest that the Ser129-p aSyn immunopositive network is a pathological phenotype, possibly representing an early stage of Lewy pathology.

**Figure 6:**
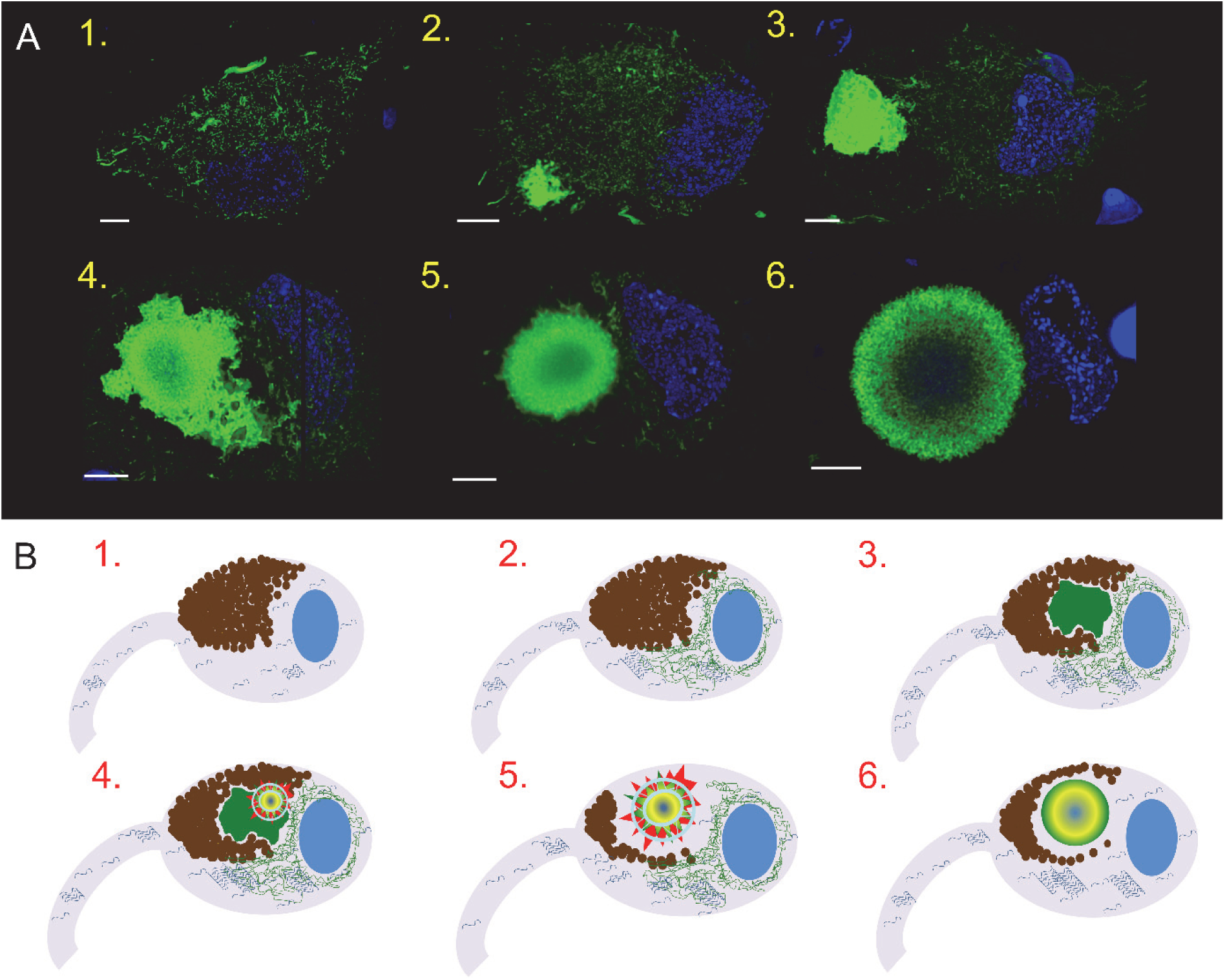
Many intracytoplasmic faces of Ser129-p aSyn hint to different maturation stages of LBs. **A:** Different patterns of Ser129-p aSyn immunoreactivity in neuromelanin-containing dopaminergic cells in the SN of PD patients, based on Ser129-p aSyn immunoreactivity patterns. Images are 2D visualizations of 3D reconstructions based on CSLM image stacks. The following cellular phenotypes were commonly observed: 1) neurons containing immunopositive intracytoplasmic network but without apparent inclusion; 2,3) neurons with intracytoplasmic immunopositive network and smaller (2) and larger (3) unstructured and expansive-appearing inclusions; 4) neuron with immunopositive intracytoplasmatic network and combination of diffusely labeled expansive-appearing inclusion and compact onion skin-type LBs; 5) neuron with intracytoplasmic network and onion skin-type LB; 6) neuron without intracytoplasmic network and onion skin-type LB. Scale bars: 5µm. **B)** Hypothetical sequence of events in LB formation in dopaminergic neurons in the SN of PD patients, based on Ser129-p aSyn immunoreactivity patterns (visualized in green; detailed explanation in text). 1) Neuromelanin-containing neuron under conditions of aSyn homeostasis; 2) alignment of Ser129-p aSyn into a cytoplasmic network; 3) sequestration of proteins into expanding inclusion; 4,5) recruitment of cytoskeletal structures to restructure the inclusion in a compaction-like manner, resulting in a multilamellar onion skin-type LB morphology (6) which reflects the encapsulation of highly aggregated proteins and lipids in its core.

### Pathological subcellular phenotypes of Ser129-p aSyn

By comparing cytoplasmic Ser129-p aSyn-immunoreactivity patterns with different specific antibodies within and between patients, we were able to identify several commonly observed reactivity patterns in melanin-containing neurons of brains with Lewy pathology. These subcellular phenotypes were strongly associated with pathology, as they were observed in patients with end-stage PD and in iLBD but not in any of the analyzed neurons in non-neurological control subjects. Representative examples are summarized in Figure 6A, as visualized by CSLM 3D reconstructions of large z-stacks of neuromelanin-containing nigral neurons in adjacent brain sections of the same patient. Commonly observed cellular phenotypes in the SN of PD patients based on Ser129-p aSyn immunoreactivity included: 1) neurons with Ser129-p aSyn immunopositive cytoplasmic network but without apparent inclusion body; 2/3) neurons with network and smaller (<5µm) or larger irregularly shaped expansive-appearing inclusions; 4) neurons revealing a Ser129-p aSyn immunopositive network and (a combination of) and uniformly stained and onion skin-like inclusions; 5) neurons with a Ser129-p aSyn immunopositive network and onion skin-type inclusions; 6) neurons without an intracytoplasmatic Ser129-p aSyn positive network, but with onion skin-type inclusion. These different faces of Ser129-p aSyn immunoreactivity possibly reflect different maturation stages of LBs (Figure 6B, as discussed later in the text), and may suggest a role for Ser129-p aSyn at different stages of Lewy inclusion formation.

## Discussion

aSyn has been established as a major component of Lewy pathology, but the subcellular distribution of specific pathology-associated forms of this protein – both within and outside the inclusions - is unclear. Detailed insight into the aSyn-based architecture of neuronal inclusions may allow for a better understaining of Lewy inclusion formation, which is a key event in PD pathophysiology. In the present study we explored the subcellular distribution patterns for CTT and Ser129-p forms of aSyn by means of specific antibody multiple labeling immunostainings in brain sections of PD patients in combination with high-resolution 3D CSLM and STED microscopy. This allowed an unprecedented detailed view on antibody-labeled subcellular pathological phenotypes, revealing several novel aspects of Lewy pathology.

First, we observed a Ser129-p-immunopositive cytoplasmic network in PD patients, using different antibodies directed against this epitope. Diffuse or granular cytoplasmic immunoreactivity has been described as a specific feature of certain antibodies against aSyn in different studies using light microscopy ^2, 29^, including antibodies with a proposed preferential affinity for disease-associated aSyn ^34, 35^. The Ser129-p aSyn cytoplasmic network was most often observed in the vicinity of a compact inclusion, although 3D revealed that this network could also be observed in neurons without apparent inclusions, indicating that this feature could represent an early phenotype of LB formation. Interesting in this perspective are the results of a previous study, which found that increased expression levels for Ser129-p aSyn in soluble fractions of cingulate and temporal cortices – as measured by western blotting – preceded the presence of histologically identified Lewy inclusions ^36^. These findings suggest a role of Ser129-p aSyn already at early stages of LB formation - thereby contradicting theories that Ser129-p aSyn occurs after LB formation ^26^ - which may have important implications for the interpretation of results of biomarker studies measuring Ser129-p aSyn in body fluids or peripheral tissues. The notion of a role of Ser129-p aSyn early in inclusion formation is supported by the experimental findings that inclusion formation after administration of recombinant (full-length and CTT) aSyn pre-formed fibrils involved the recruitment of soluble endogenous aSyn and its intracellular phosphorylation at Ser129 ^37, 38^. Possible roles of Ser129-p at this stage could be a stabilizing effect on aggregating proteins ^39^ - Ser129-p was demonstrated to inhibit aSyn fibrillogenesis *in vitro* ^26, 40^ - while Ser129-p aSyn has also been suggested to serve as an activator of autophagic instruments ^41, 42^.

The different localization of the Syn105 and 11A5 antibodies in midbrain LBs and dystrophic LNs, directed against 122CTT and Ser129-p aSyn, respectively, has been described before^21^. Here, we confirm this finding using new highly-specific antibodies against the same epitopes (Supplementary Table 5 and Supplementary Figures 9 and 10), and for different CTT aSyn (calpain-cleaved at res. 119 and 122) species. We further observed a separation between antibodies against NT/NAC domains and CT domains of aSyn in LBs. Integrating antibodies selectively directed against the above mentioned aSyn PTMs and domains in multiple labeling protocols revealed a 3D onion skin-like orchestration of LBs, with concentric lamellar bands enriched for specific aSyn epitopes. The consistency of this lamination was revealed by semi-quantitative analysis in a subset of nigral LBs and different PD patients. The multilamellar appearance of LBs has been described in different studies that focused on the ultrastructure of LBs using EM techniques ^43, 44^, while lamination patterns in LBs and dystrophic LNs were also suggested in studies using light microscopy ^21, 45, 46^. Interestingly, in the present study this lamellar phenotype was visualized by the gradual distribution of immunoreactivities for antibodies directed against different epitopes on one single protein, aSyn.

We explored whether the differences in the distribution of aSyn epitopes throughout nigral LBs were related to the distribution of proteins and lipids in these structures using CARS microscopy, a label-free imaging technique. Unfortunately, due to the impaired morphological integrity of the fresh-frozen tissue sections after CARS imaging, a detailed subclassification of LB morphologies by high-resolution CSLM or STED imaging was not possible. However, the results provided interesting insights into the biochemical composition of aSyn immunopositive inclusions. First of all, we observed substantial heterogeneity in the content of proteins and lipids between inclusions within subjects. Results showed that increased lipid content could be detected in a fraction of nigral Ser129-p positive inclusions. In this subset of lipid-enriched inclusions, lipids were found to be centralized in the core of this structure, together with increased protein content. In a previous study, we demonstrated the presence of lipid and membraneous structures in LBs using correlative light and electron microscopy (CLEM) and TEM, which was confirmed by CARS and Fourier transform infrared spectroscopy (FTIR) ^32^. Moreover, reactivity in the core of LBs has further been described for different lipophilic dyes ^32, 46, 47^. The functional relevance behind clustering of lipids in the center of LBs is not clear at this point. However, extensive experimental evidence has demonstrated the binding of aSyn to (bio)membranes, while the presence of lipid molecules was repeatedly reported to increase the aggregation rate of aSyn ^48^.

At the periphery of nigral onion skin-type LBs, the band of Ser129-p aSyn immunoreactivity was embedded in an organized cage-like framework of cytoskeletal components, including beta-tubulin and intermediate neurofilaments. The structured presence of cytoskeletal components, major constituents in organizing cellular organelles and substructures, at the periphery of LBs is suggestive of the active encapsulation of proteins and lipids in the core of these structures. This theory was supported by the results of label-free CARS microscopy, which showed increased protein and lipid content mainly at the core of aSyn-positive inclusions. The displacement of intermediate neurofilaments from their normal cellular distribution and their encapsulation of aggregated proteins have been previously described as consistent features of intracellular aggresome formation ^49, 50^. Although the arrangement of neurofilament at the rim of Lewy-type pathology has been reported before ^31^, this study provides important new detailed insights into the structured organization of such components at the LB periphery. Thereby, our results confirm previous studies proposing that LBs share phenotypic features with aggresomes ^51^. Our observations indicate that the interplay of Ser129-p aSyn with cytoskeletal proteins may be an important step in the process of LB morphogenesis.

Based on Ser129-p Syn immunoreactivity patterns within and between PD patients, we identified a subset of commonly observed subcellular pathological phenotypes in neuromelanin-containing neurons in the SN of PD patients (Figure 6A), which may reflect different maturation stages of LB pathology. Based on our high-resolution observations in different experimental setups, combined with CARS data, we propose a hypothetical sequence of events in the formation of LBs in the SN (Figure 6B). We speculate that different subcellular phenotypes of aSyn pathology are tightly coupled to the progressive collapse of protein degradation systems. In healthy dopaminergic neurons, the basal proteolytic activity of the intracellular protein degradation systems – in particular the ubiquitin-proteasomal system (UPS) and chaperone-mediated autophagy (CMA)-are able to maintain protein homeostasis (Figure 6B, step 1) ^52^. In situations of increased protein burden, these systems are overloaded, and superfluous aSyn could start to aggregate with itself, other proteins, membranes and/or organelles ^32^. Extensive phosphorylation of aSyn may take place to stabilize the expanding mass of cellular debris and to activate the macrophagic machinery ^42^. The Ser129-p aSyn-immunopositive intracytoplasmic network may reflect the alignment and sequestration of aggregated material for focused clearance by means of autophagy and aggrephagy (step 2) ^49, 50^. When the expansive-appearing inclusion cannot be not cleared (step 3), it will continue to grow and occupy an increasing surface in the cytoplasm. At a certain point, cytoskeletal systems may be recruited to the LB to actively restructure the inclusion into a compact and stable morphology (step 4, 5). Interestingly, the idea of a restructuring of LBs during their maturation in a compaction-like manner has been proposed before in literature ^2^. In the mature onion skin-like morphology, highly aggregated proteins and lipids are centralized in the core of the structure, encapsulated by a cage-like framework of Ser129-p aSyn and cytoskeletal components (step 5, 6). This hypothetical sequence of events in LB maturation could be further explored in future experimental studies, for instance in cellular and animal models of aSyn aggregation in PD.

In line with previous studies using antibodies against CTT aSyn species, we found two CTT variants to localize towards the core of LBs ^17, 21, 53^. Moreover, the different localization between antibodies against residues 118-126 of aSyn’s CT and NT/NAC antibodies suggested that a large part of aSyn in the LB core has a shortened CT. In addition, this may suggest that other CTT variants than 119CTT and 122CTT also localize preferntially in the LB core. CTT of aSyn was repeatedly found to increase the propensity of aSyn to form amyloid aggregates *in vitro* ^16, 22, 23, 25^, and these observations together have led to the hypothesis that CTT aSyn plays a critical role in the initiation of protein aggregation ^21^. However, as LBs were demonstrated to contain a medley of fragmented membranes and organelles ^32^, including components that are able to cleave aSyn, for instance caspase-1 ^25^, it cannot be ruled out that the enrichment of CTT aSyn in the core of LBs and LNs is the result of post-aggregation events. Interestingly, the results of a recent study in a neuronal aSyn PFF model suggested that the post-aggregation CTT of aSyn (primarily at Glu-114) plays a key role in the cellular processing of aSyn fibrils^38^. Amongst others, CTT of aSyn led to efficient packaging of aSyn fibrils in inclusion bodies^38^. These results support an extensive cellular regulation of Lewy inclusion formation and a crucial role for aSyn PTMs in this process.

Punctate cytoplasmic reactivity for 122CTT aSyn only showed limited co-localization with the Ser129-p immunoreactive network, and was independent of the presence of Lewy pathology (Supplementary Figure 6A). This PTM has been previously suggested to be a normal cellular process ^16^ and, indeed, presence of CTT aSyn was detected in brains of non-neurological control subjects by western blotting ^17^. Interestingly, a substantial part of the 122CTT aSyn immunoreactive punctae in the cytoplasm was observed to localize at the outer membrane of VDAC/Porin-reactive mitochondria in diseased neurons from PD patients, but also in healthy neurons from non-neurological control subjects. This could be placed in line with the findings of the previously mentioned study, in which a 15kDa band corresponding with CTT aSyn, was observed in fractions enriched for lysosomes and mitochondria derived from SH-SY5Y cells expressing human WT α-synuclein ^17^. Future experimental studies are necessary to explore the functional relevance of the co-occurence of 122CTT aSyn and mitochondria.

The immunoreactivity of aSyn antibodies often surrounded an unstained central core, which showed converging immunoreactivity for DAPI - a dye that binds to T-A-rich regions of DNA ^54^. This feature has already been reported before ^55^. Although at this point the relevance of this observation is not clear, it was speculated that this may be the result of mitochondrial DNA incorporated in LBs ^55^. Alternatively, DAPI may interact with certain aggregated proteins or lipids in the center of LBs. The limited immunoreactivity in the core of LBs may be the result of limited accessibility of antigen in this densely packed domain ^46^ or of masking or destruction of epitopes by its strong chemical environment. Importantly, LBs were shown to contain many (>300) proteins as identified by proteomics, which are predominantly centralized at the core of LBs based on our CARS results.

In summary, the present study provides a STED perspective on the architecture of Lewy pathology. Our results reveal a structured onion skin-like distribution of different forms of aSyn in nigral LBs and LNs. Our data suggest that LBs are actively regulated, structured encapsulations of aggregated proteins and lipids by a cage-like framework of Ser129-p aSyn and cytoskeletal components. Analysis of subcellular reactivity patterns led to the identification of different pathology-associated, Ser129-p aSyn-immunopositive subcellular phenotypes, suggesting a central role for Ser129-p aSyn in Lewy inclusion formation. The applied combination of well-characterized highly specific antibodies and super-resolution microscopy techniques in this study allowed an unprecedented detailed phenotyping of antibody-labeled subcellular pathology, which opens exciting opportunities for better characterization and understanding of LB formation in the pathology of PD.

## Material and methods

### Postmortem human brain tissue

Postmortem human brain tissue from clinically diagnosed and neuropathologically verified donors with advanced PD as well as non-demented controls was collected by the Netherlands Brain Bank (www.brainbank.nl). In compliance with all local ethical and legal guidelines ^56^, informed consent for brain autopsy and the use of brain tissue and clinical information for scientific research was given by either the donor or the next of kin. The procedures of the Netherlands Brain Bank (Amsterdam, The Netherlands) were approved by the Institutional Review Board and Medical Ethical Board (METC) from the VU University Medical Center (VUmc), Amsterdam. Brains were dissected in compliance with standard operating protocols of the Netherlands Brain Bank and Brain Net Europe.

The details of all donors included in this study are listed in Supplementary Table 2. Most of these PD donors developed symptoms of dementia during their disease course (Supplementary Table 2), and had extensive α-synuclein pathology throughout the brain (Braak LB stage 5/6) ^33^. In addition, donors with earlier Braak stages (Braak LB stage 3/4) were included in our study, as well as iLBD cases that did not develop clinical Parkinson’s disease but displayed Lewy pathology in their brain (Braak LB stage 3) ^33^. Formalin-fixed paraffin-embedded (FFPE) tissue blocks of the substantia nigra (SN) and hippocampus - also containing part of the parahippocampal gyrus- from these donors with PD, iLBD and also 6 non-demented controls (details in Supplementary Table 2) were cut into 10 and 20 µm thick sections, which were utilized for immunohistochemistry and multiple labeling experiments. In addition, fresh-frozen tissue blocks of the SN from 5 patients with advanced PD were cut into 10 µm for CARS microscopy (specified in Supplementary Table 2).

### Generation and initial characterization of aSyn antibodies

A detailed overview of all utilized antibodies and their epitopes in this study is provided in Supplementary Table 1. Generation and characterization of antibody 11A5 and syn105 was previously described ^14, 57, 58^. Additional novel antibodies were generated by immunizing rabbits either with E. coli derived recombinant full length aggregated aSyn (asyn-055 and asyn-058) or KLH-conjugated peptides representing the C-terminus of 119CTT, 122CTT aSyn, or aSyn derived peptide phosphorylated at Ser129, respectively (for asyn-131, asyn-134, asyn-142; Supplementary Table 5 and Supplementary Figures 9 and 10. After screening of serum titers, standard B cell cloning was performed to generate rabbit monoclonal antibodies (mAbs). Recombinant mAbs were screened for binding to the peptides representing the aa1-60, 61-95, and aa96-140 by ELISA, respectively (for asyn-055 and asyn-058), or the C-terminus of aSyn119CTT, aSyn122CTT or phosphorylated at Ser129, respectively (for asyn-131, asyn-134, asyn-142), by ELISA and surface plasmon resonance (SPR). For asyn-055 and asyn-058, counter-screen by ELISA was performed with beta- and gamma-synuclein. For asyn-131, and asyn-134 ELISA- or SPR-based counter-screenings using C-terminal elongated peptides were performed to identify mAbs highly specific for the C-termini of 119CTT or 122CTT aSyn. ELISA- or SPR based counter-screenings using the corresponding non-phosphorylated peptide were performed to identify asyn-142 as highly specific for aSyn phosphorylated at Ser129. All animal experiments followed highest animal welfare standards and were performed according to ethics protocols approved by the local animal welfare committee at Roche, while animal experiment licenses were approved by the respective state authorities. For all novel antibodies used in this study, details on their characterization are provided in Supplementary Table 5 and Supplementary Figure 9. Additionally, in Supplementary Figure 10, specificity of all aSyn mAbs used in this study is shown based on Western blot with different recombinant forms of aSyn.

Antibodies included in the multiple labeling experiments were first optimized for immunohistochemistry. In the multiple labeling for the studied aSyn epitope, immunoreactivity patterns were validated using at least one different antibody raised against a similar epitope, with exception of the antibodies against 119CTT and the N-terminus of aSyn, as no other available antibody against a similar epitope could be integrated in the multiple labeling protocol.

### Immunohistochemistry

Protocols for the antibodies against aSyn were optimized for light microscopy to characterize their immunoreactivity in human postmortem formalin-fixed paraffin-embedded brain tissue. All IHC protocols could be optimized without antigen retrieval procedure and without addition of Triton. The EnvisionTM+ kit (DAKO, Santa Clara, USA) was used as a high-sensitivity visualization system, with 3,3’-diaminobenzidine (DAB; 1:50 diluted in substrate buffer; DAKO) as the visible chromogen. Stained sections were analyzed using a Leica DM5000 B photo microscope (Leica Microsystems, Heidelberg, Germany). All brightfield images included in Supplementary Figure 1 were acquired using a HC PL APO 63×1.40 oil objective using a Leica DFC450 digital camera (Leica Microsystems).

### Development of triple and multiple labeling protocols

#### 1. Immunoreactivity patterns of aSyn epitopes

Using immunofluorescent stainings, antibodies against different domains and PTMs of aSyn were co-visualized and their local distribution patterns were assessed in pathological structures and neurons. Triple labeling experiments, including DAPI and two antibodies against aSyn were performed to obtain insight into their distribution patterns. Moreover, to allow systematic comparison of distribution patterns of different aSyn epitopes, protocols were developed to visualize multiple (4 or 5) antibodies against aSyn in the same section. To validate findings from the initial multiple labeling experiments, different antibodies against similar epitopes were selected. This ‘validation set’ of antibodies was optimized for additional multiple labeling protocols. The sets of antibodies used in the different protocols are specified in Supplementary Table 1. No antigen retrieval methods or permeabilization steps were applied in any of these experiments.

For each protocol, we made use of a combination of direct and indirect immunodetection methods. Several primary antibodies (specified in Supplementary Table 1) were directly labeled with fluorochromes following standard protocols of different labeling kits (art. no. A20181, A20183, A20186, 21335 for labeling with Alexa 488, Alexa 546, Alexa 647, and biotin, respectively; Thermo Fisher Scientific, Waltham, USA). Each protocol started with an indirect immunolabeling using unlabeled primary antibodies raised in rabbit/mouse using their appropriate secondary antibodies (with different conjugates, specified in Supplementary Table 1). Sections were then blocked for 1 hour in 5% normal rabbit serum and 5% normal mouse serum in PBS. After this, a biotinylated primary antibody (raised in mouse or rabbit) could be incubated, and visualized by fluophore-conjugated streptavidin. Then, sections were incubated in blocking solution (2% normal goat serum) containing the diluted directly labeled antibodies together with DAPI (1 µg/ml). Sections were mounted in Mowiol mounting solution using glass cover slips (Art. No.: 630-2746; Glaswarenfabrik Karl Hecht, Sondheim, Germany). Negative control stainings lacking primary antibodies were performed to control for background/autofluorescence levels and aspecific staining. Single stainings using a directly labeled antibody against Ser129-p aSyn were scanned to determine autofluorescence levels of the studied morphological structures (LBs, LNs), which was found negligible under the applied scan settings.

#### Association CTT and Ser129-p aSyn with subcellular markers

In order to study the association of immunoreactivity of CTT and Ser129-p aSyn with subcellular structures, additional multiple labeling protocols were further designed. Apart from the described antibodies against aSyn, these protocols also included some commercial antibodies as markers for subcellular structures, including mitochondria, ER and cytoskeletal proteins (Supplementary Table 1). In these protocols, heat-induced epitope retrieval using citrate buffer (pH 6) and a permeabilization step (1hr incubation in 0.1% Triton-x) was included. Negative control stainings lacking primary antibodies were included to control for background/autofluorescence levels and aspecific staining.

### Confocal and STED microscopy

CSLM and STED microscopy were performed using a Leica TCS SP8 STED 3X microscope (Leica Microsystems). All images were acquired using a HC PL APO CS2 100× 1.4 NA oil objective lens, with the resolution set to a pixel size of 20 nm x 20 nm. All signals were detected using gated hybrid detectors in counting mode. Sections were sequentially scanned for each fluorophore, by irradiation with a pulsed white light laser at different wavelengths (indicated in Supplementary Table 1). Stacks in the Z-direction were made for each image. To obtain CSLM images of the DAPI signal, sections were irradiated with a solid state laser at a wavelength of 405 nm. For STED imaging, a pulsed STED laser line at a wavelength of 775 nm was used to deplete Abberior (580, 635P), Alexa (594, 647) or Li-Cor (680 nm) fluorophores, while continuous wave (CW) STED lasers with wavelengths of 660 nm and 592 nm were used to deplete the Alexa 546 and Alexa 488 fluorophores, respectively. The DAPI signal was not depleted, so this channel was scanned at the same resolution as the CSLM images.

After scanning, deconvolution was performed using CMLE (for CSLM images) and GMLE algorithms in Huygens Professional (Scientific Volume Imaging; Huygens, The Netherlands) software. Images were adjusted for brightness/contrast in ImageJ (National Institute of Health, USA). 3D reconstructions were made using the LAS X 3D Visualization package (Leica Microsystems). Final figures were composed using Adobe Photoshop (CS6, Adobe Systems Incorporated).

### Image processing and semi-quantitative analysis

Nigral LBs were classified and selected for inclusion in the analysis based on their immunopositivity for Ser129-p in combination with morphological criteria (specified in Results section). Additional criteria for inclusion were 1) the diameter of the structure (at least 5µm) and 2) the presence of specific signal for all channels (signal intensity of raw CSLM images substantially higher than autofluorescence or background levels under the applied scan settings). In this selected subset of LBs, distribution patterns of immunoreactivities were analyzed on deconvolved CSLM images of 30 LBs in the SN of 8 patients with advanced PD (Supplementary Table 2). Z-stacks were made for each structure, of which three frames in the central portion of the structure (Z length: 0.30µm; step size between frames: 0.15 µm) were selected to quantify the x-y distribution for different markers. For the analysis, a maximum Z-projection of these selected frames was first made in ImageJ. Subsequently, three 100 px (2µm) thick lines were drawn over three equatorial planes of the LBs (similar to ^46^) in ImageJ, along which signal intensities for each channel were measured using a script. The average intensity for each channel at each point of the diameter was normalized to its maximum intensity in the same structure, while the position along the diameter was expressed as % diameter. Normalized values were used to generate average line profiles per morphological structure. The center of the LB was defined as the origin of the structure ^46^. The position in the LB with the maximum intensity was determined per channel. Ranking of absolute positions of maximum intensities per structure with respect to the origin of the LB were compared between channels (nonparametric Friedman test). P-values for multiple comparisons were adjusted using Dunn’s correction for multiple comparisons. Statistical analyses were done using SPSS software (version 22, IBM) and GraphPad software (version 7.0, Prism).

### Coherent anti-Stokes Raman scattering

The workflow used for CARS microscopy is outlined in Supplementary Figure 11. The detection of the lipid and protein distribution was performed on native, dried samples ^59, 60^. A commercial setup (Leica TCS SP5 II CARS, Leica Microsystems) was used with an HCX IRAPO L25X/0.95W (Leica Microsystems) objective. For the lipid distribution intensity images were taken at 2850 cm^−1^ (Pump-wavelength 816 nm, Stokes-wavelength 1064 nm) and for the protein distribution intensity images at 2930 cm^−1^ (Pump-wavelength 810 nm, Stokes-wavelength 1064 nm). The laser power at the sample was 28 mW (Pump) and 21 mW (Stokes). Integration times of 34 s per image with a pixel dwell time of 32 µs, 1024×1024 pixels and a spatial resolution of 300 nm were used ^61^. After the label-free detection of the lipid and protein distribution, immunofluorescent stainings were performed on the same sections (Supplementary Figure 11). Tissue sections were fixed in 4% formaldehyde for 10 minutes and stained for aSyn, using two primary antibodies raised against aSyn (LB509; ab27766, Abcam, Cambridge, UK) and Ser129-p aSyn (ab59264, Abcam) and their appropriate secondary antibodies. After this, sections were incubated in Sudan Black for 30 min and mounted in Mowiol. For fluorescence detection, a commercial setup (Leica TCS SP5 II CARS, Leica Microsystems, Heidelberg, Germany) was used. Data evaluation was done in Matlab with the Image Processing and Statistics toolboxes (The Mathworks, Inc., Mass.,USA). First, large overview CARS-intensity and fluorescence images were manually overlaid by comparison of morphological features. The distribution of aSyn, proteins and lipids was identified by the overlay of both fluorescence images (Supplementary Figure 11). Therewith, autofluorescence of the surrounding tissue and the fluorescent signal of the aSyn-immunopositive inclusions could be separated. The inclusion bodies were manually identified based on morphology. Only inclusions with a diameter of 5-20 µm were included for analysis.

### Image processing and analysis of CARS images

For an objective evaluation of CARS intensity of aSyn-immunopositive inclusion bodies, the mean CARS intensity of the direct surrounding, a donut with a width of 3.5 µm, was compared with the CARS intensity of the inclusion (Supplementary Figure 10A, light blue and yellow area). The areas of aSyn-immunopositivity were transferred into the CARS-intensity images. Areas with no intensity in the CARS intensity images (holes) were excluded by intensity thresholding. CARS-pixel-intensities higher than 1.4 times the mean CARS-intensity of the surrounding were defined as higher protein/lipid content, which was determined based on pilot measurements in a subset of (∼40) aSyn-positive inclusions. The ratio between the CARS-pixel-intensities of the LB and the mean CARS intensity of the surrounding were calculated and the areas with higher protein/lipid content were marked in red (Supplementary Figure 12). Morphological filtering and image processing were performed in Matlab R2017a, MathWorks.

## Code availability

Custom-written ImageJ scripts are available upon request.

## Data availability

All images and data supporting the findings of this study are available upon request.

## Acknowledgements

We are grateful to all individuals that donated their brain to the Netherlands Brain Bank (NBB; www.brainbank.nl). We thank the team of the NBB, in particular Michiel Kooreman, for their cooperation and their help in the selection of brain tissue. We thank the Advanced Optical Microscopy Core O|2 (www.ao2m.amsterdam) for support with STED imaging. Further, we thank Lidia Janota for performing immunofluorescent stainings of the tissue sections used for CARS microscopy.

## Author contributions

T.M. and C.M. performed immunohistochemistry and multiple labeling experiments. T.M., C.M., E.T., J.K. and W.vdB. performed STED imaging, as well as processing and analysis of images. D.N., D.P., S.EM. and K.G. performed experiments and data analysis for CARS microscopy. D.M. performed the labeling of antibodies for multiple labeling experiments and contributed to the experimental design. M.B., L.S., W.Z., R.B., O.M., K.K., S.H., M.H., T.K., M.R., S.D. selected and characterized the aSyn antibodies. T.M., W.vdB, J.G.and M.B. designed research, analyzed and interpreted the data, and contributed to writing the manuscript.

---

**Supplementary Table 1:** Antibodies used in final multiple labeling experiments with direct and indirect detection. Exp: experiment. 1) light microscopy; 2) multiple labeling of aSyn epitopes; 3) multiple labeling of aSyn epitopes with CTT119; 4) multiple labeling of aSyn epitiopes, validation set; 5) multiple labeling PTMs and cytoskeletal markers; 6) multiple labeling PTMs + ER; 7) multiple labeling PTMs and mitochondria. *: biotinylated antibody used in multiple labeling protocols. When PTM-specific is indicated in the Table, the antibody was raised to specifically recognize PTM protein variants of aSyn, while other antibodies were designed to recognize specific sequence motifs in the C-terminus. Further details on the specificity of these antibodies for different forms of aSyn are presented in Supplementary Table 5 and Supplementary Figures 9 and 10. **Abbreviation:** AB ID: antibody identifier; Exp: experiment; Ref: reference for additional information on AB characterization; d.l.: antibody directly labeled with fluophore.

| Epitope | AB ID (Cat #) | Source | Host species | Exp | Fluorochrome | STED depletion wavelength | Ref |
| --- | --- | --- | --- | --- | --- | --- | --- |
| Ser129-p (PTM-soecicic) | 11A5 | Prothena | Mouse | 1,2,5,6,7 | Alexa 488(d.l.)/Alexa 488 | 592 | <sup>14</sup> |
| Ser129-p (PTM-specific) | asyn-142 | Roche | Rabbit | 1,4,5,6,7 | Alexa 488 (d.l.)/Alexa 488 | 592 |  |
| 122CTT (PTM-specific) | Syn105 | Prothena | Rabbit | 1,2*,5*,6*,7 | Alexa 547<br>Alexa 594/Abberior 635p | 660<br>775 | <sup>57</sup> |
| 122CTT (PTM-specific) | asyn-134 | Roche | Rabbit | 1,4*,5*,6*,7 | Alexa 547<br>Alexa 594/Abberior 635p | 660<br>775 |  |
| 119CTT (PTM-specific) | asyn-131 | Roche | Rabbit | 1,3,5,7 | Alexa 647 (d.l.) | 775 |  |
| NT (40-55) | 23E8 | Prothena | Mouse | 1,4 | Alexa 647 (d.l.) | 775 |  |
| NAC | asyn-055 | Roche | Rabbit | 1,2 | Alexa 488<br>Li-Cor 680 | 592<br>775 |  |
| NAC | asyn-058 | Roche | Rabbit | 1,4 | Li-Cor 680 | 775 |  |
| CT (118-126) | 5C1 | Prothena | Mouse | 1,2 | Alexa 594 | 775 | <sup>58</sup> |
| CT (121-125) | SC-211 | Santa Cruz | Mouse | 1,4 | Alexa 594 | 775 |  |
| KM-51 | MONX10738 | Monosan | Mouse | 1 | - (only light microscopy) | - |  |
| Intermediate neurofilament | 837801 | BioLegend | Mouse | 5,6 | Abberior 580 | 775 |  |
| Beta-tubulin | ab18207 | Abcam | Rabbit | 5 | Abberior 635p | 775 |  |
| Calreticulin | ab2907 | Abcam | Rabbit | 6 | Abberior 635p | 775 |  |
| VDAC/Porin | ab14734 | Abcam | Mouse | 7 | Abberior 580 | 775 |  |

**Supplementary Table 2:** Brain tissue of different donors included in the experiments. **Abbreviations:** CD: clinical diagnosis; CERAD: CERAD age-related score for neuritic plaques; Exp: Experiments: 1) FFPE sections used for immunofluorescence multiple labeling experiments; 2) Snap-frozen sections used for CARS microscopy; Three of these donors (indicated by *) were also studied in the study by Shahmoradian et al. (Ref 42). The identifier of these donors in this study is added to the table. PDD: PD with dementia; AD: Alzheimer’s Disease

| ID | Diag. | Age at death | Sex | Braak aSyn stage | Braak NFT stage | CERAD score | Age of diagnosis | Age at onset dementia | Exp | Figure | Donor ID in REF42 |
| --- | --- | --- | --- | --- | --- | --- | --- | --- | --- | --- | --- |
| PD1 | PDD | 80 | M | 6 | 1 | B | 55 | 73 | 1 | 2A,E, S5 |  |
| PD2* | PDD | 72 | M | 6 | 1 | O | 64 | 69 | 1 | S1A | **J** |
| PD3 | PD | 72 | M | 6 | 1 | A | 57 | - | 1 | - |  |
| PD4* | PDD | 80 | M | 6 | 1 | O | 69 | 77 | 1 | S1B;<br>4B,D;<br>S3A,C, S5 | **L** |
| PD5* | PD | 77 | M | 6 | 2 | O | 59 | - | 1 | 2B,C | **A** |
| PD6 | PD+AD | 74 | M | 6 | 4 | B | 64 | 73 | 1 | 1B;<br>4A,B,D;<br>5A,C; S5,<br>S6 |  |
| PD7 | PDD | 66 | F | 6 | 1 | A | 57 | 64 | 1 | 1A; 2D;<br>4C,E;5E;<br>S3D;<br>S2,S4 |  |
| PD8 | PDD | 87 | M | 6 | 3 | C | 77 | 86 | 1 | 1C |  |
| PD9 | PD | 67 | F | 6 | 1 | B | 60 | - | 2 | - |  |
| PD10 | PDD | 81 | M | 6 | 2 | O | 72 | 79 | 2 | 3C |  |
| PD11 | PDD | 74 | M | 5 | 3 | O | 67 | 71 | 2 | 3D |  |
| PD12 | PDD | 76 | M | 6 | 1 | B | 66 | 71 | 2 | 3A, B |  |
| PD13 | PDD | 83 | M | 6 | 1 | B | 78 | 80 | 1,2 | S5 |  |
| PD14 | PD | 90 | F | 3 | 2 | A | 85 | - | 1 | S8 |  |
| PD15 | PDD | 85 | F | 4 | 2 | C | 74 | 82 | 1 | - |  |
| iLBD1 | C | 93 | F | 3 | 2 | O | - | - | 1 | S8 |  |
| iLBD2 | C | 86 | F | 3 | 3 | A | - | - | 1 | S8 |  |
| iLBD3 | C | 98 | M | 3 | 2 | O | - | - | 1 | S8 |  |
| C1 | C | 83 | F | 0 | 3 | A | - | - | 1 | S6A |  |
| C2 | C | 80 | M | 0 | 1 | B | - | - | 1 | - |  |
| C3 | C | 89 | F | 0 | 1 | A | - | - | 1 | S6B |  |
| C4 | C | 84 | F | 0 | 1 | O | - | - | 1 | - |  |
| C5 | C | 81 | M | 0 | 2 | O | - | - | 1 | - |  |
| C6 | C | 70 | F | 0 | 2 | A | - | - | 1 | - |  |

**Supplementary Table 3:**
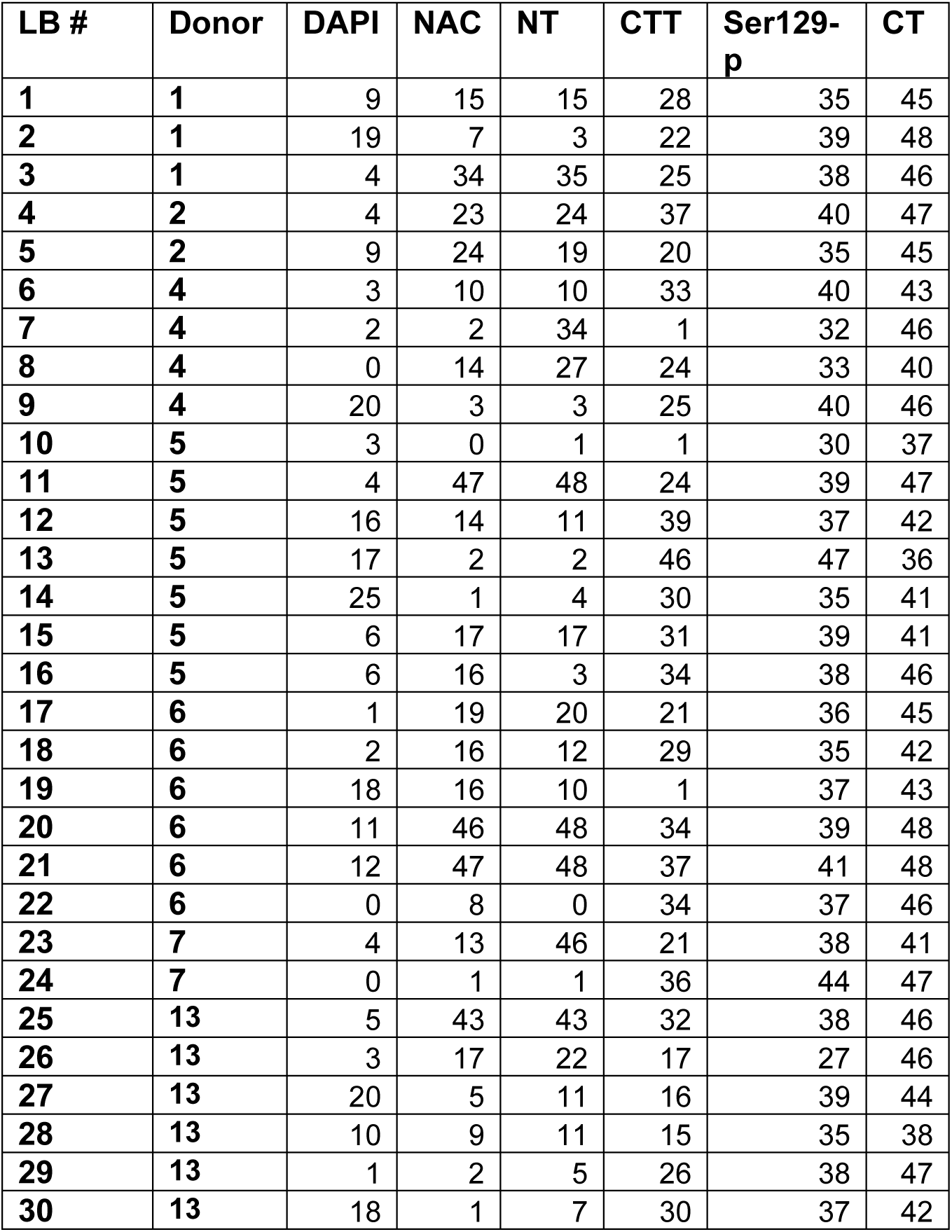
Number of structures scanned per patient and locations for peak intensities for different aSyn epitopes in the semi-quantitative measurement of line profiles. Values represent the distance of peak intensities relative to the origin of the LB (with values of 0 representing the LB origin, 50 the peripheral border, Figure 2F).

**Supplementary Table 4:** Different localization of peak intensities between aSyn epitopes. Overview of Dunn’s corrected p-values of multiple comparisons. Ns = not significant; *: p<0.05; **p<0.01; ***p<0.001

|  | NAC | NT | 122CTT | Ser129-p | CT |
| --- | --- | --- | --- | --- | --- |
| NAC | X | 1,00 | <0.001*** | <0.001*** | <0.001*** |
| NT | ns | X | ns | 0.008** | <0.001*** |
| 122CTT | <0.001*** | ns | X | 0.025* | <0.001*** |
| Ser129-p | <0.001*** | 0.008** | 0.025* | X | 0.05* |
| CT | <0.001*** | <0.001*** | <0.001*** | 0.05* | X |

**Supplementary Table 5.** Specifications of novel aSyn mAbs generated at Roche. Specific binding of aSyn Mabs was assessed by ELISA, custom-made PepStarTM peptide microarrays (JPT, Berlin, Germany), or surface plasmon resonance (SPR). For ELISA, the full length human recombinant αSyn, βSyn or αSyn proteins, indicated fragments or peptides were coated or linked by biotin tag on an ELISA plate, incubated with primary antibody and primary antibody was detected by anti-rabbit secondary antibody with a colorimetric readout (OD, optical density). For the peptide array, peptide sequences had a length of 15 amino acid residues and were designed to cover the whole sequence of human aSyn. Neighboring peptides had an overlapping sequence of 11 amino acids. Primary antibodies were hybridized with the array and detected by fluorescently labelled anti-rabbit IgG antibodies following the manufacturers’ instruction. For SPR see Supplementary Figure 9. Binding late indicates the binding strength of the peptide towards the mAb at the end of the association phase. aa, amino acids; CTT, C-terminal truncation; ELISA, enzyme-linked immunosorbent assay; hu, human; NAC, non-β-amyloid component of Alzheimer’s plaques region of aSyn; rec., recombinant; RU, relative unit.

Antibody specifications – Roche antibody set 1
| Epitope | mAb | hu rec. aSyn Monomer Binding EC50 (ng/ml)<br>hu rec. aSyn, aa1-60 (OD)<br>hu rec. aSyn, aa96-140 (OD) | Protein array for full length α-, β-, or γSyn<br>Peptide array-based epitope mapping, immunoreactivity for indicated peptides<br>Overall conclusion |
| --- | --- | --- | --- |
| NAC | asyn-055 (rabbit) | 1508.0<br>0.0<br>0.0 | Binds to αSyn but not βSyn or γSyn<br>aa65-79 (weak), aa93-107, aa97-111 (weak)<br>Based on ELISA and epitope mapping, presumable conformational epitope in NAC region |
| NAC | asyn-058 (rabbit) | 1167.7<br>0.0<br>0.0 | No binding to αSyn, βSyn, γSyn<br>No detection of immunoreactivity to any peptide<br>Based on clear result in ELISA, presumable conformational epitope in NAC region |

Antibody specifications – Roche antibody set 2
| Epitope | mAb | OD on ELISA / SPR binding late (RU) in kinetic screening mode |
| --- | --- | --- |
| aSyn 119CTT | asyn-131 (rabbit) | aSyn(106-119): 1.119 / 324<br>aSyn(106-122): 0.088 / 14 |
| aSyn 122CTT | asyn-134 (rabbit) | aSyn(111-122): 1.368 / 523<br>aSyn(111-124): 0.075 / 1 |
| aSyn pSer129 | asyn-142 (rabbit) | aSyn(122-135, pSer129): 1.209 / 152<br>aSyn(122-135): 0.145 / 3 |

**Supplementary Figure 1:**
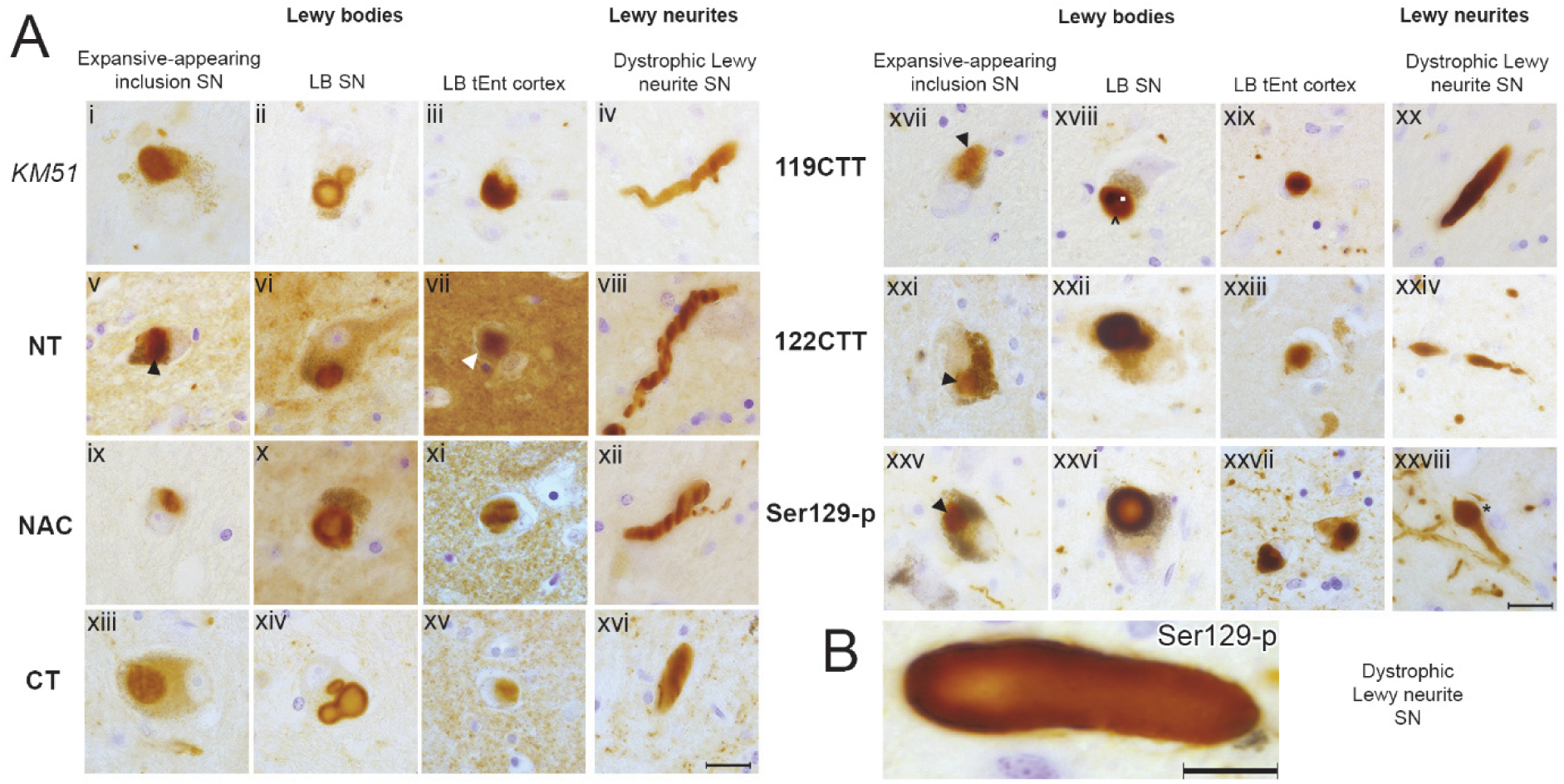
Characterization of immunoreactivity for antibodies against aSyn domains and PTMs in postmortem brains with advanced PD pathology. **A:** Representative brightfield images for selected morphological structures detected by antibodies against different aSyn epitopes. All images were taken in the SN or transentorhinal cortex of patient PD2. Different IHC features are flagged (discussed in text). Black and white arrowheads highlight aSyn-immunopositive neuronal inclusions. In xviii, a weaker ring of immunopositivity (white dot) surround the portion with strongest labeling (indicated with a ^). In xxviii, a dystrophic LN is indicated by an asterix. B: Example of a dystrophic LN with with stronger immunolabeling at its periphery compared to its core, as observed using an antibody against Ser129-p aSyn (11A5) in the SN of patient PD4. Scale bar = 10µm. C: Schematic outline of the aSyn protein together with the epitopes for the different antibodies used in the present study (See also Supplementary Table 1). A: Scale bar = 20µm; B: scale bar = 10µm.

**Supplementary Figure 2:**
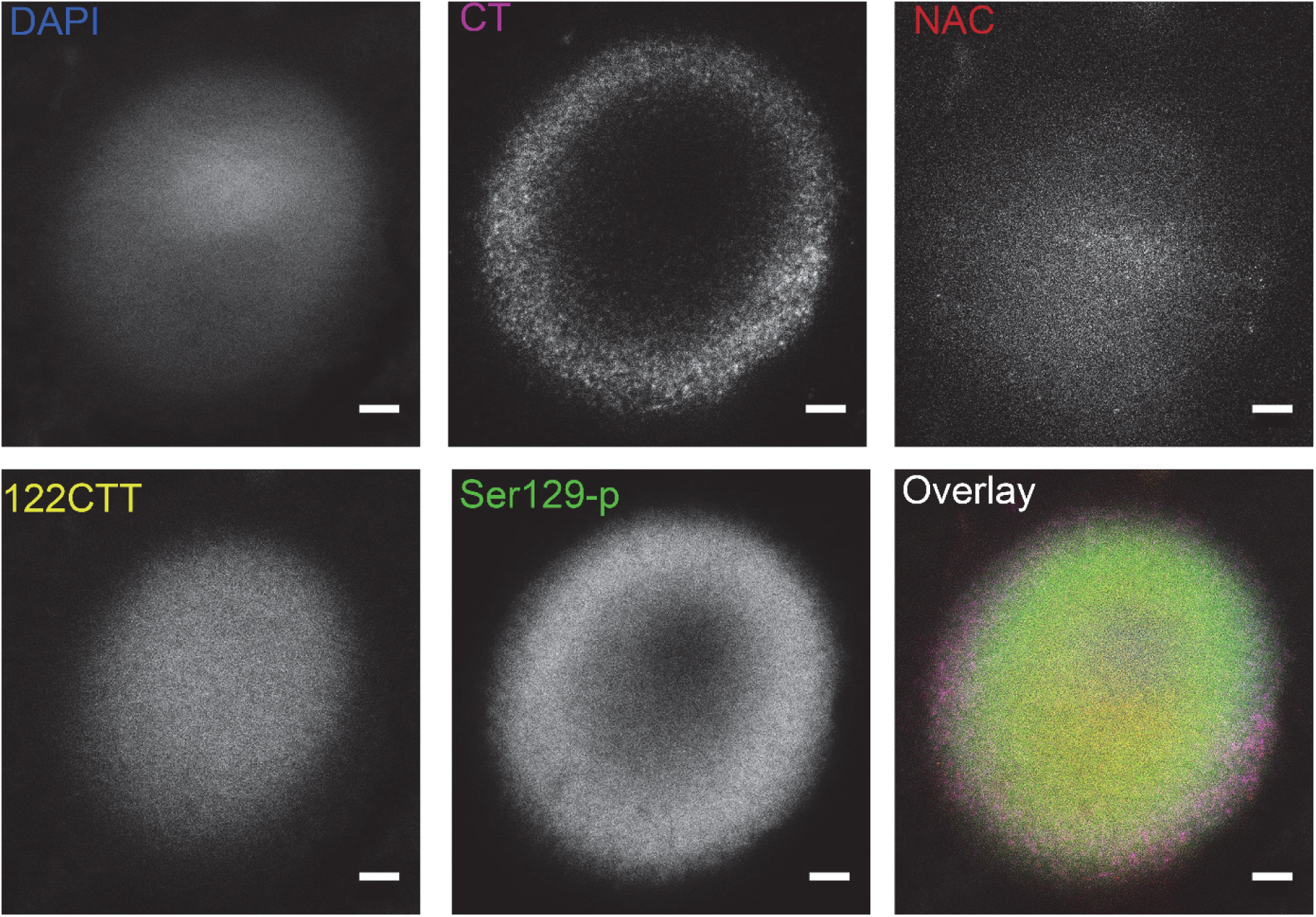
Multilamellar pattern of immunoreactivities for different aSyn epitopes in onion-skin like LBs is observed with different antibodies (validation set). Raw STED images taken in the SN of patient PD7. Scale bar: 2µm

**Supplementary Figure 3:**
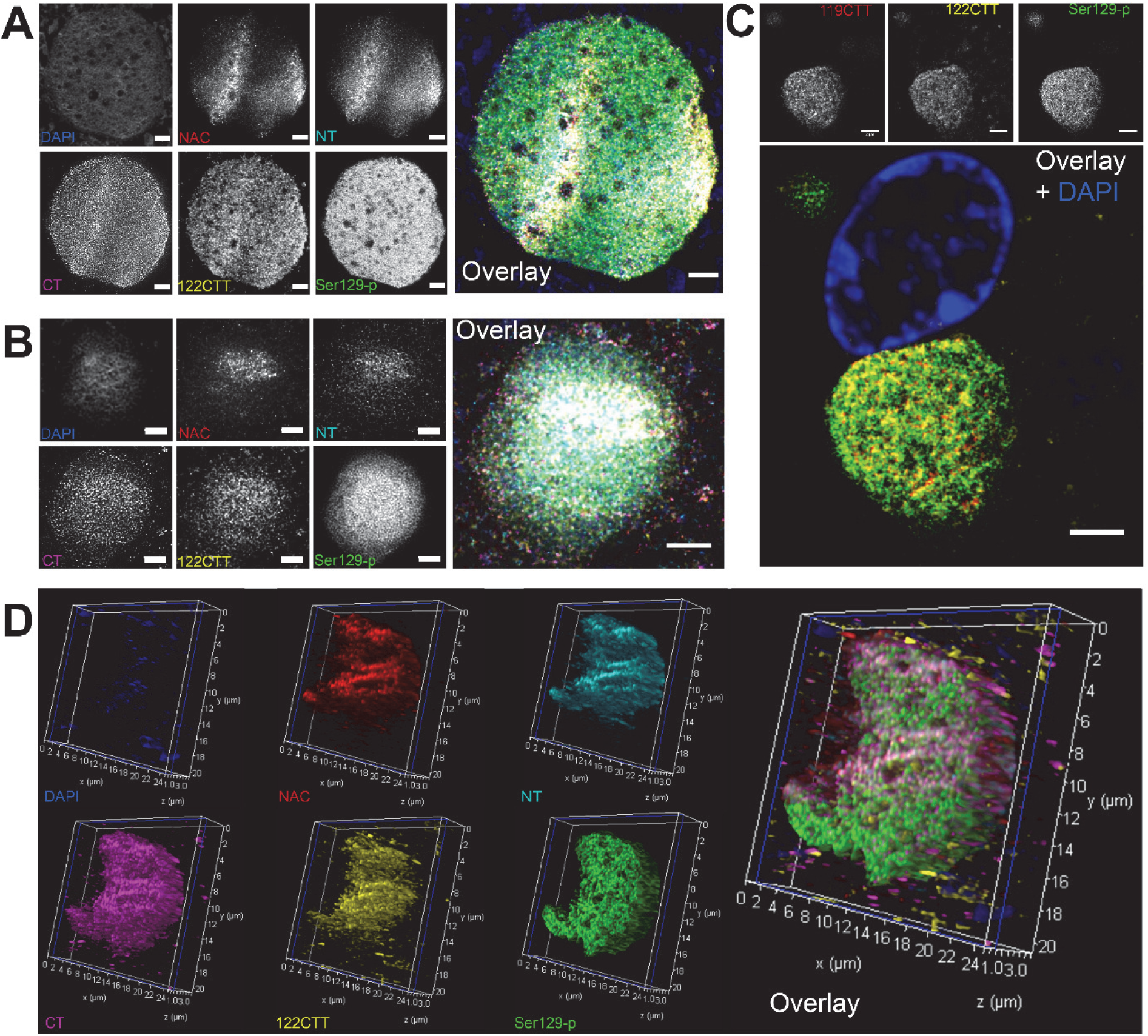
Distribution patterns of aSyn PTMs and domains in nigral expansive-appearing aSyn inclusions and cortical LBs. **A. B. C:** Representative deconvolved STED images of nigral expansive-appearing inclusions in the SN, and LBs in the transentorhinal cortex. and hippocampus of patient PD4. The distribution of immunoreactivities was generally more diffuse, although CT and Ser129-p aSyn immunoreactivity at the edges of the structure often sharply delineated the inclusion. D. 3D reconstruction of a expansive-appearing inclusion body in the SN based on deconvolved CSLM images, taken in patient PD7. A-C: Scale bar = 2 µm

**Supplementary Figure 4:**
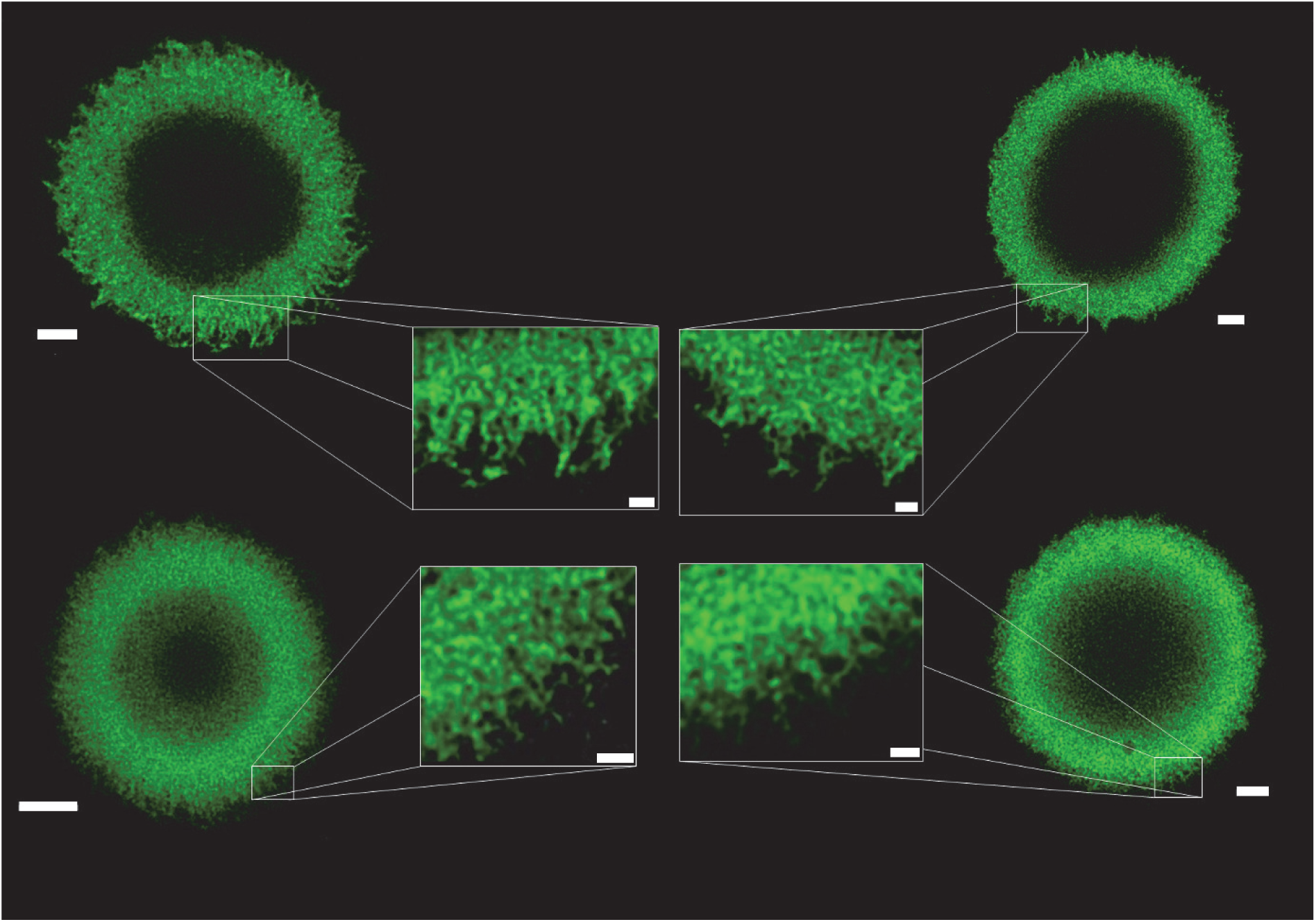
Gallery of radiating Ser129-p immunoreactivity patterns at the periphery of nigral LBs. STED deconvolved images taken in the SN of different PD patients. Scale bars main Figure = 2μm; scale bars insets = 0.5μm

**Supplementary Figure 5:**
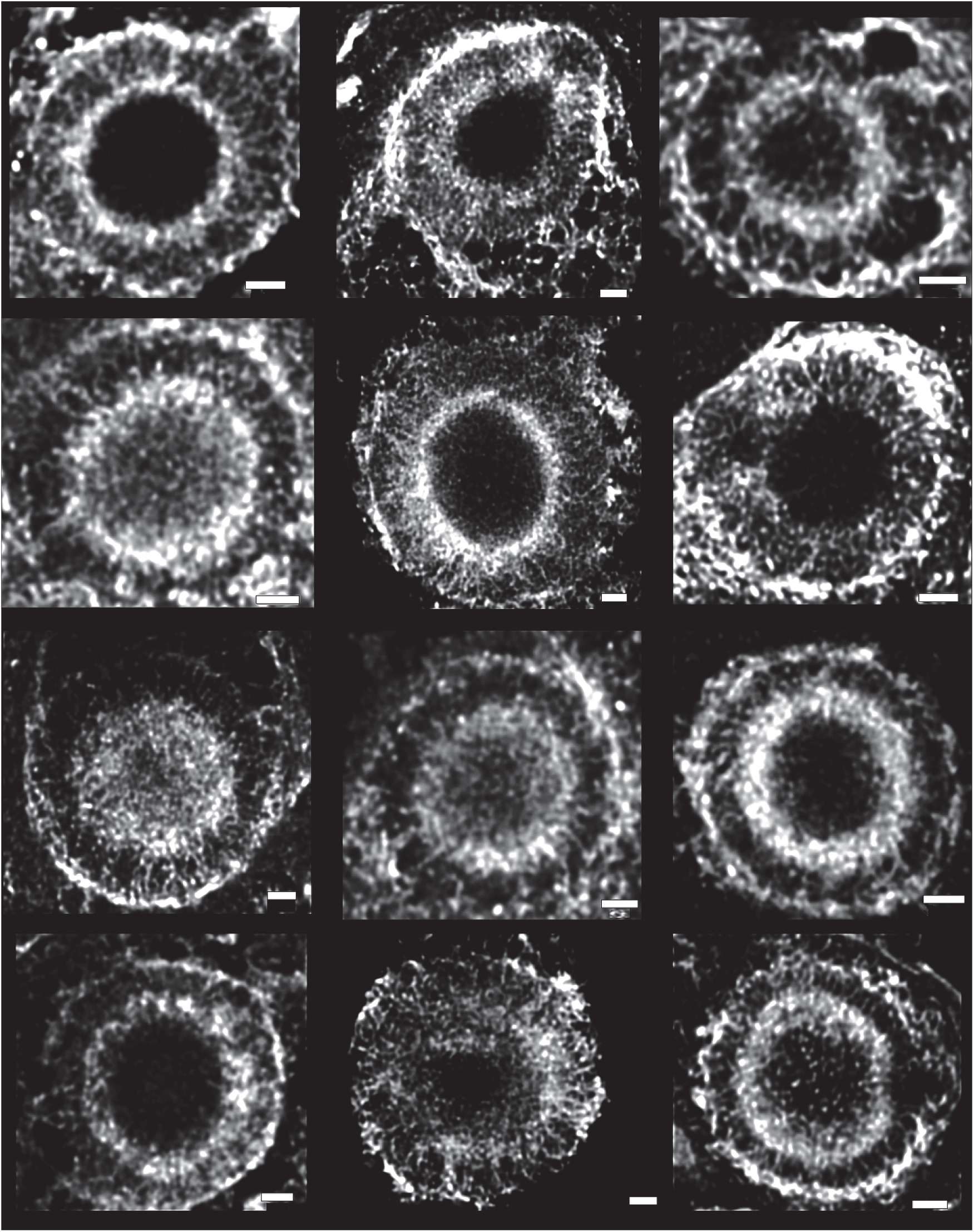
Gallery of ‘wheel-like’ neurofilament structures in nigral LBs immunopositive for Ser129-p aSyn (not shown). Deconvolved CSLM images captured in the SN of various patients (indicated in Supplementary Table 2). Scale bars = 2μm.

**Supplementary Figure 6:**
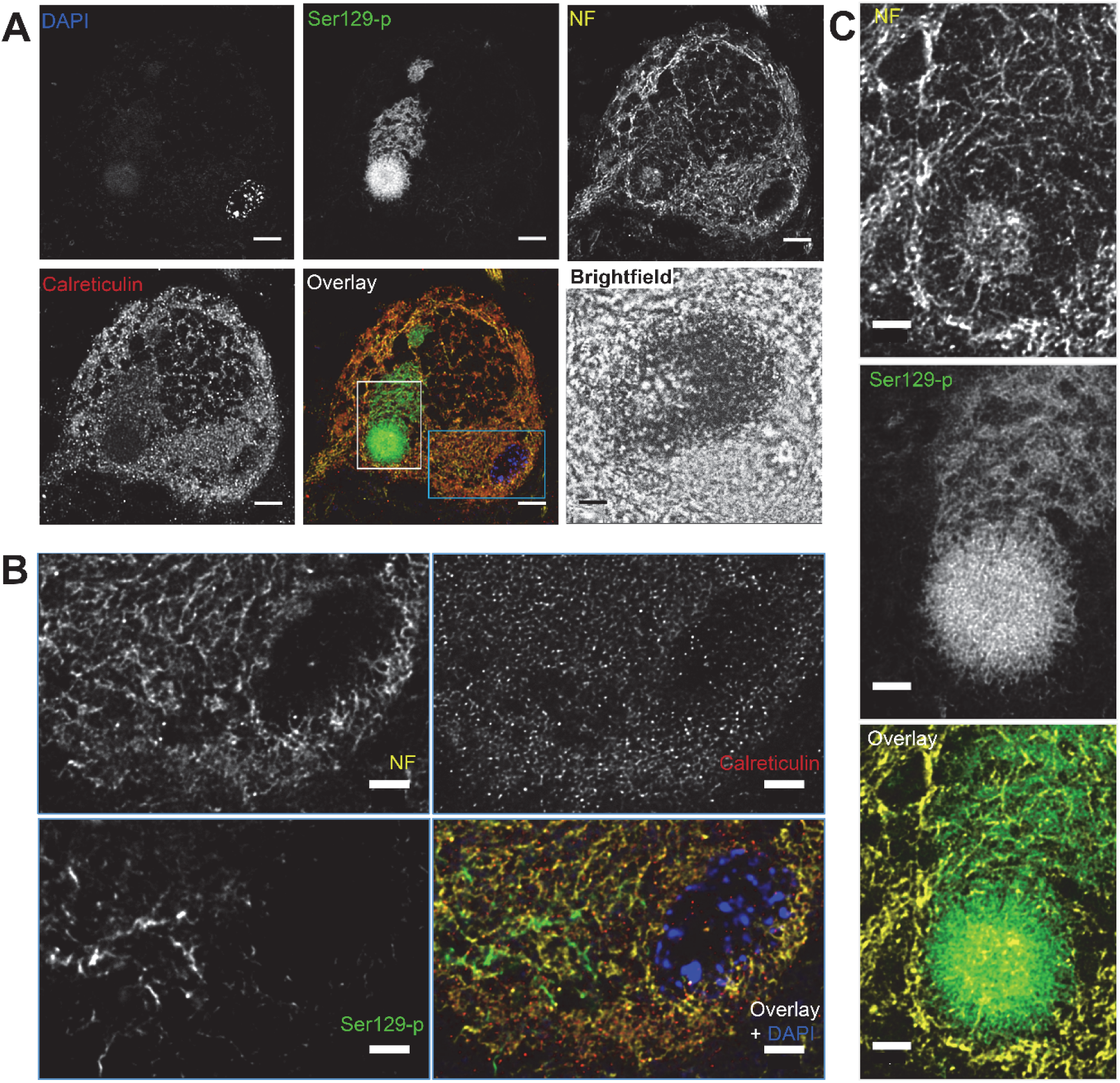
Intracytoplasmic 129Ser-p aSyn network shows limited co-localization with other intracytoplasmatic networks. Images were taken in a dopaminergic neuron the SN of a PD patient containing a combination of a combination of expansive appearing inclusion and a small LB with a barely distinguishable ring-structure. A: Overview: deconvolved CSLM images of the entire neuron showing different intracytoplasmatic networks reactive for calreticulin (ER marker), neurofilament and Ser129-p aSyn. Note the limited availability of epitope in the area containing neuromelanin (visible in brightfield image as dark material). B: Detailed view on the area in the blue square indicated in A. Deconvolved STED images showing limited co-localization between markers for intracytoplasmic networks of neurofilament. C: Detailed view of the inclusion (area indicated in the grey square in A). The compact-appearing spherical LBs shows 1) radiating immunoreactivity patterns for Ser129-p aSyn; 2) a cage-like framework of neurofilament. NF: intermediate neurofilament. A: Scale bar = 5µm; B, C: scale bar = 2 µm

**Supplementary Figure 7:**
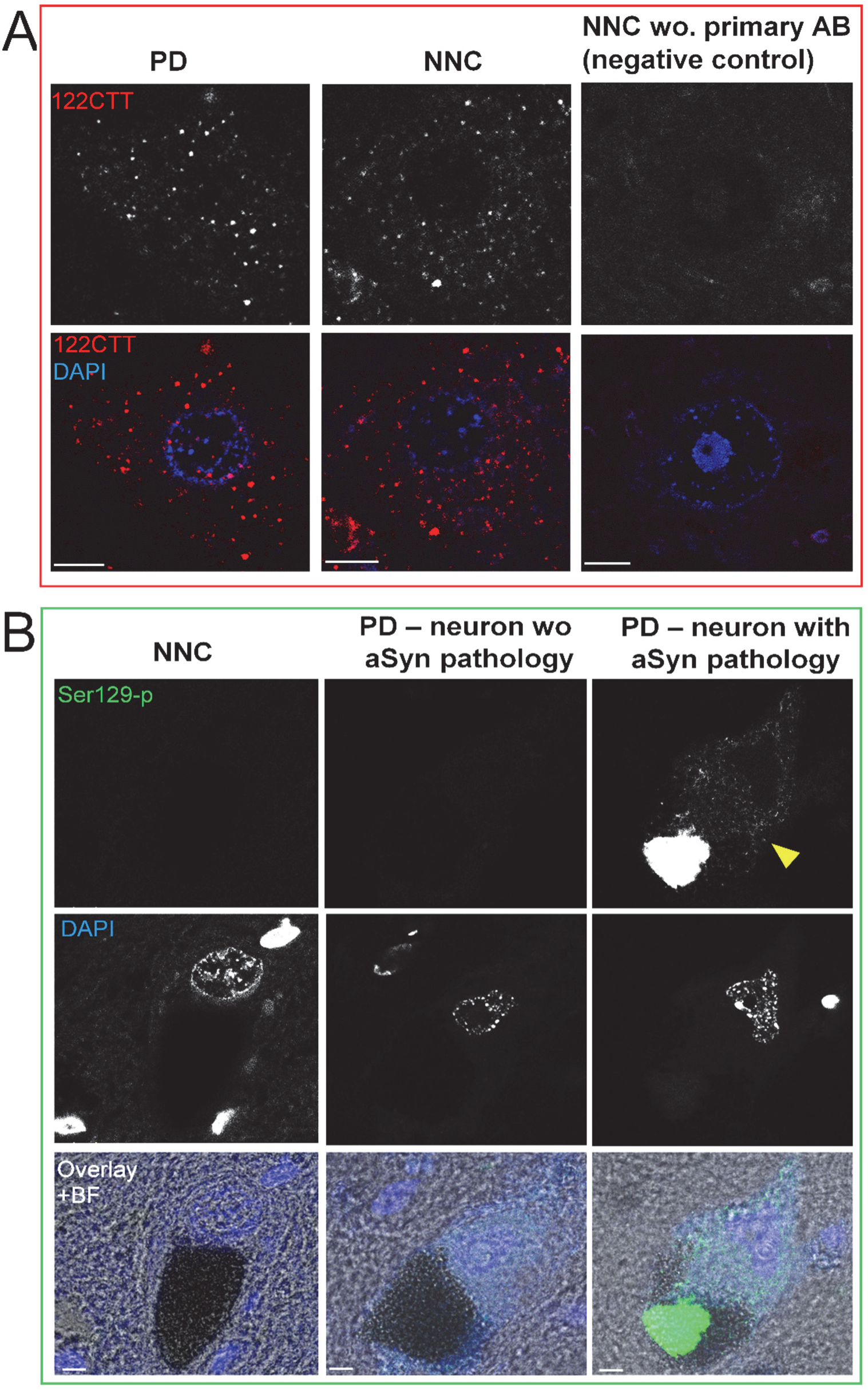
Subcellular reactivity patterns for CTT and Ser129-p aSyn in brains without Lewy pathology. **A:** The dot-like pattern of 122CTT aSyn was visible both in PD patients and brains without Lewy pathology (NNC = non-neurological control). However this pattern was not observed in negative controls lacking the primary antibody. Images taken in the hippocampus. B: The cytoplasmic Ser129-p positive network (indicated by yellow arrowhead) was only observed in certain cells in PD patients – often associated with the presence of an inclusion - and not in non-demented controls. Raw CSLM images that were scanned, processed and visualized in the same way. Images taken in melanin-containing cells in the SN. A,B: Scale bar = 5µm. Abbreviations: NNC = non-neurological control; BF: brightfield.

**Supplementary Figure 8:**
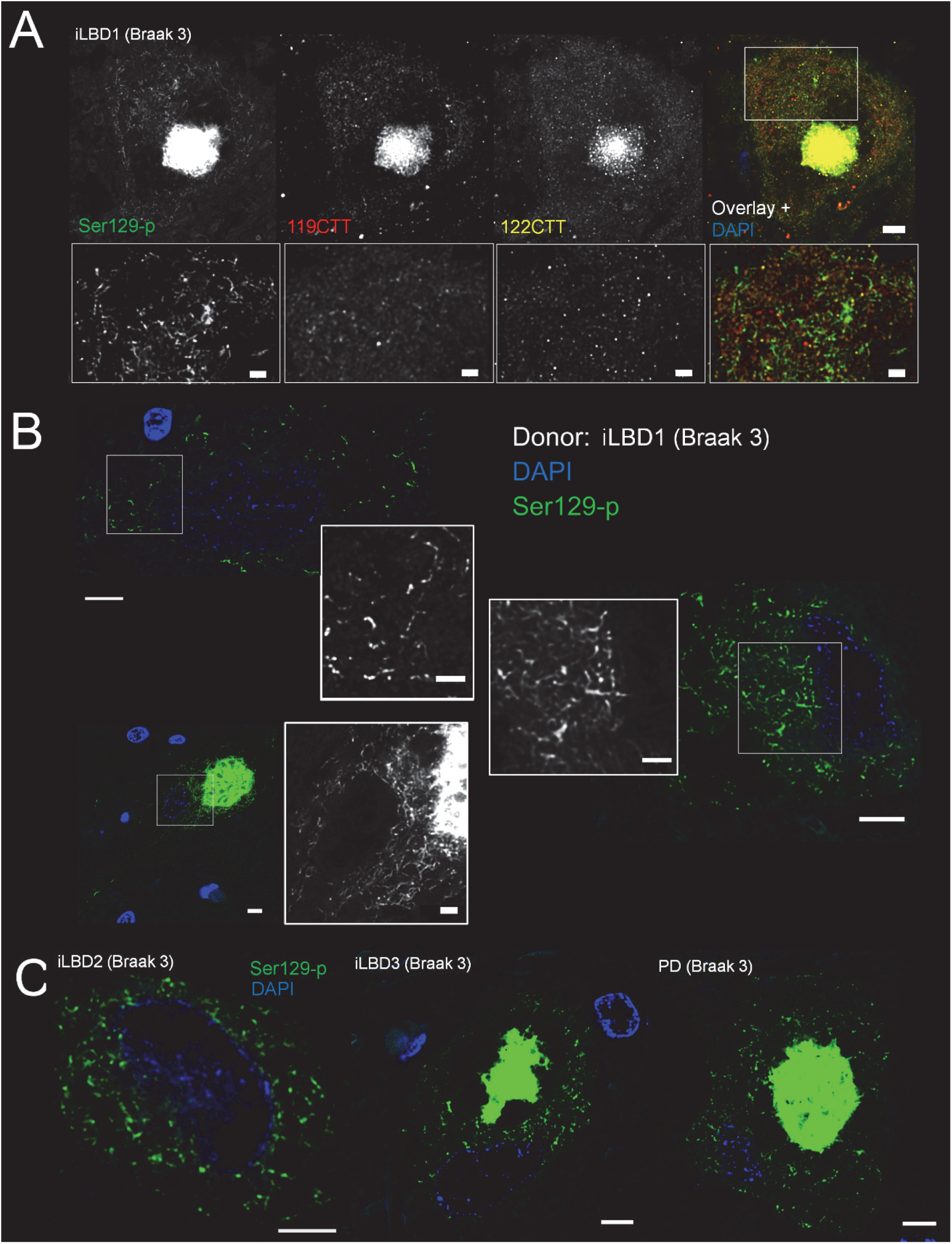
PTM aSyn immunoreactivity patterns in donors with iLBD and PD with Braak stage 3 for LBs. **A:** Ser129-p aSyn immunopositive network is present in the SN of donors with iLBD, but is not highlighted by antibodies against 119CTT or 122CTT aSyn – despite of strong co-localization of these PTMs in LBs. B: More detailed view on some of the Ser129-p aSyn immunopositive cytoplasmic networks in neuromelanin-containing dopaminergic neurons with and without apparent inclusions. Please note that some of these features were very subtle in cells without inclusion (top left) as opposed to the dense network that is visualized in neurons with expansive-appearing inclusion bodies. C: Examples of Ser129-p aSyn immunoreactive networks in other iLBD and PD patients (indicated in Supplementary Table 2). Deconvolved CSLM images. A: Upper row: scale bar = 5 µm; lower row: scale bar = 2 µm; B: main image: scale bar = 5 µm; inset: scale bar = 2 µm; C: scale bar = 5 µm.

**Supplementary Figure 9.**
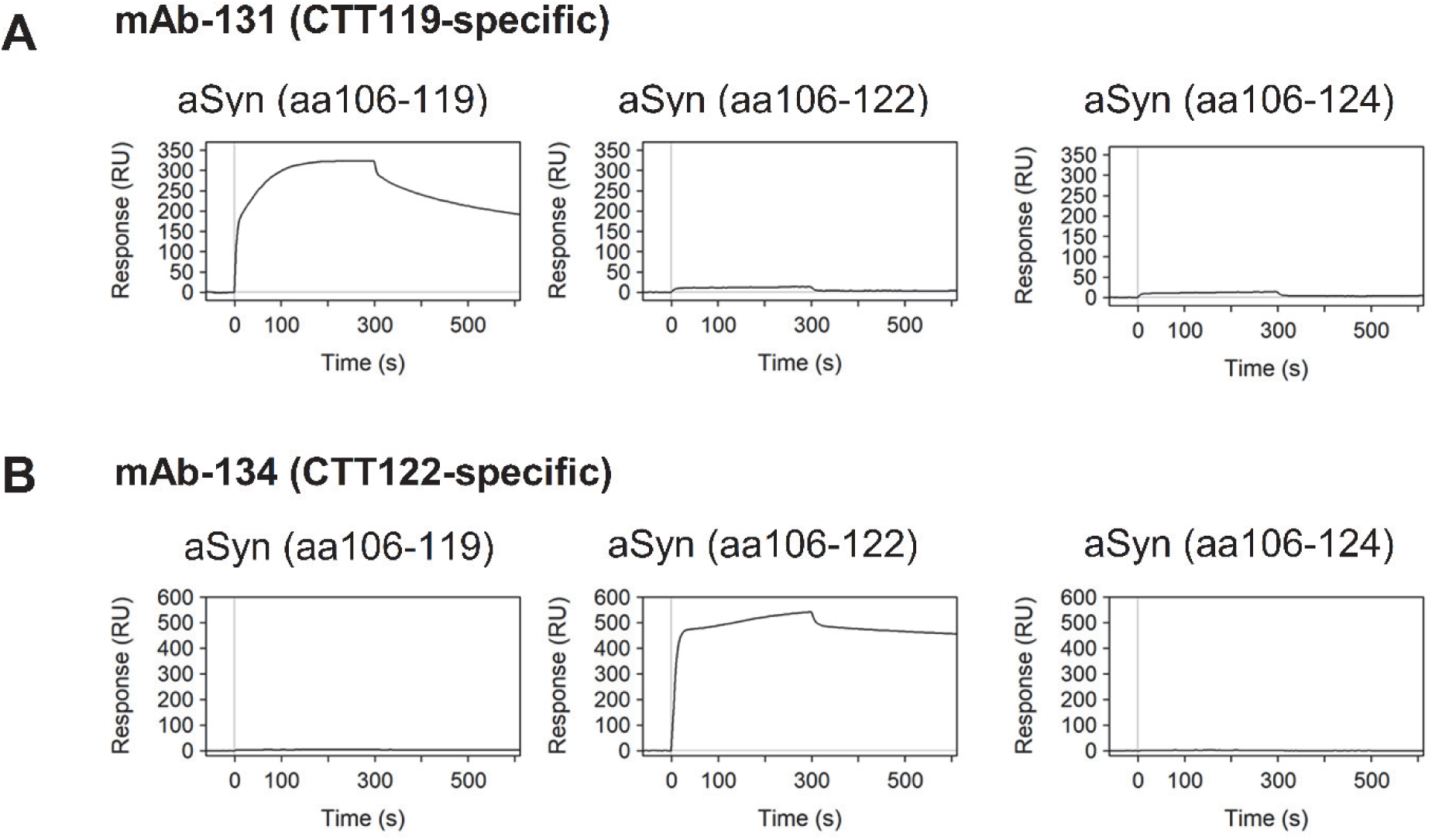
Specificity of novel Roche mAbs directed against truncated or phosphorylated α-synuclein. The specificity of asyn-131mAb (A) and asyn-134 mAb (B) against truncated isoforms of α-synuclein has been analyzed by surface plasmon resonance (SPR) using the mAbs captured by a Fc specific anti-rabbit polyclonal antibody on the sensor chip and biotinylated peptides corresponding to the C-termini of truncated α-synuclein (i.e. aSyn(aa 106-119) or aSyn(aa 106-122), respectively) as analytes. Biotinylated C-terminal elongated peptides (aSyn(aa 106-122) or aSyn(aa 106-124), respectively) have been used as negative control to demonstrate the specificity of the mAbs for the truncated C-terminal sequence. The asyn-142 mAb against α-synuclein phosphorylated at Serine-129 (C) was analyzed similarly, whereas biotinylated peptides corresponding to Serine-129 phosphorylated or non-phosphorylated α-synuclein (i.e. aSyn(aa 122-135, pSer129) or aSyn(aa 122-135), respectively) were employed as analytes. Biotinylated peptides have been grafted with streptavidin to increase the sensitivity of the SPR analysis. Analyses have been performed on a Biacore B4000 instrument at 25°C according to a method described by Schräml and Biehl.^62^

**Supplementary Figure 10:**
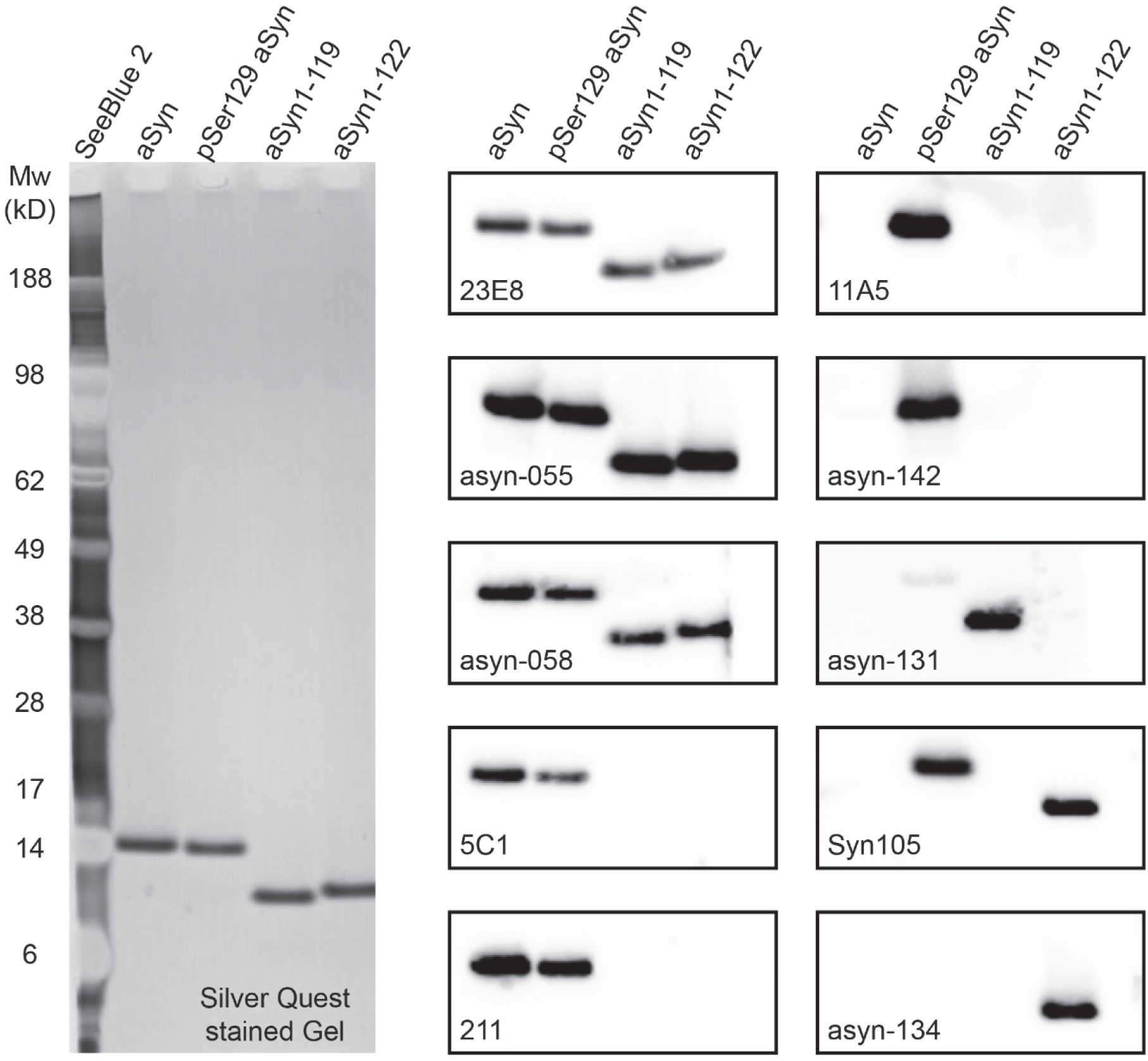
Specificity of various aSyn mAbs based on Western blot with different recombinant forms of aSyn. ***Method*:** Recombinant forms of full length aSyn, pSer129 aSyn, aSyn1-119, and aSyn1-122 were expressed tag-free in E. coli and purified by well-established protocols. 10 ng of protein was run per lane on a 4-12% Bis/Tris polyacrylamide gel under denaturing conditions and transferred to a nitrocellulose membrane for Western blotting. One gel was stained by Silver Quest to visualize total protein load. Western blots were blocked by Super Block (Thermo 37515) followed by 5% milk, each for 15min at RT and then incubated with 0.01 μg/ml (syn105), 0.1 μg/ml (211), or 1μg/ml (all others) primary antibody concentration and bound antibody was detected with anti-mouse IgG-HRP (211, 23E8, 5C1, 11A5) or anti-rabbit IgG-HRP (all others). Images of optimal exposure times (1-18seconds) of individual blots are shown. *Results*: As expected, 23E8 (epitope region within aa40-55), asyn-055, and asyn-058 (both with epitopes within the NAC region) bind all forms of analyzed recombinant aSyn. Binding of both C-terminal aSyn Mabs 5C1 (epitope in aa118-126) and 211 (epitope within aa121-125) was abolished in aSyn1-119 and aSyn1-122 whereas full length aSyn and pSer129 aSyn were recognized as expected, respectively. This confirms that these antibodies are highly sensitive to aSyn cleavage at aa119 and aa122 but are not influenced by pSer129 posttranslational modification. Note the strong specificity of 11A5 and asyn-142 for pSer129aSyn (both mAbs were generated and selected for binding pSer129 aSyn), of asyn-131 to aSyn1-119 (generated and selected to bind aSyn truncated at aa119), and of asyn-134 to aSyn1-122 (generated and selected to bind aSyn truncated at aa122) and the complete lack of biding to other forms of aSyn by all these antibodies, confirming their high specificity and selectivity. Syn105 (polyclonal rabbit antibody generated by immunization toward the peptide CGGVDPDN which contains the aa118-122 sequence of aSyn, VDPDN) was previously reported to have some concentration-dependent cross-reactivity towards full length recombinant human aSyn in ELISA-based characterization and on Western blots with extracts of brain tissue from human aSyn transgenic mice and patients with Dementia with Lewy bodies^20^. Similar to what was demonstrated in Games et al. by ELISA, Syn105 showed on our Western blot at 0.01 μg/ml strong immunoreactivity towards aSyn1-122 and it was not able to detected full length aSyn and neither did it bind to aSyn1-119. In contrast, we observed for this lot of Syn105 on Western blot an unexpected immunoreactivity towards recombinant pSer129 aSyn. It is unclear whether this an in vitro artifact that translates to binding of pSer129 aSyn expressed in human brain during immunofluorescence analysis as performed in this paper. In previous immunohistochemical analysis of brain sections from human aSyn transgenic mice and patients with Dementia with Lewy bodies, Games et al. demonstrated that co-localization of pSer129 immunoreactivity (polyclonal rabbit IgG^20^, Millipore) with immunoreactivity of Syn105 was limited. Nevertheless, to address the potential issue of cross-reactivity of Syn105 in our studies, we employed the highly 122CTT-specific and selective mAb asyn-134 to validate the STED-findings obtained by Syn105.

**Supplementary Figure 11:**
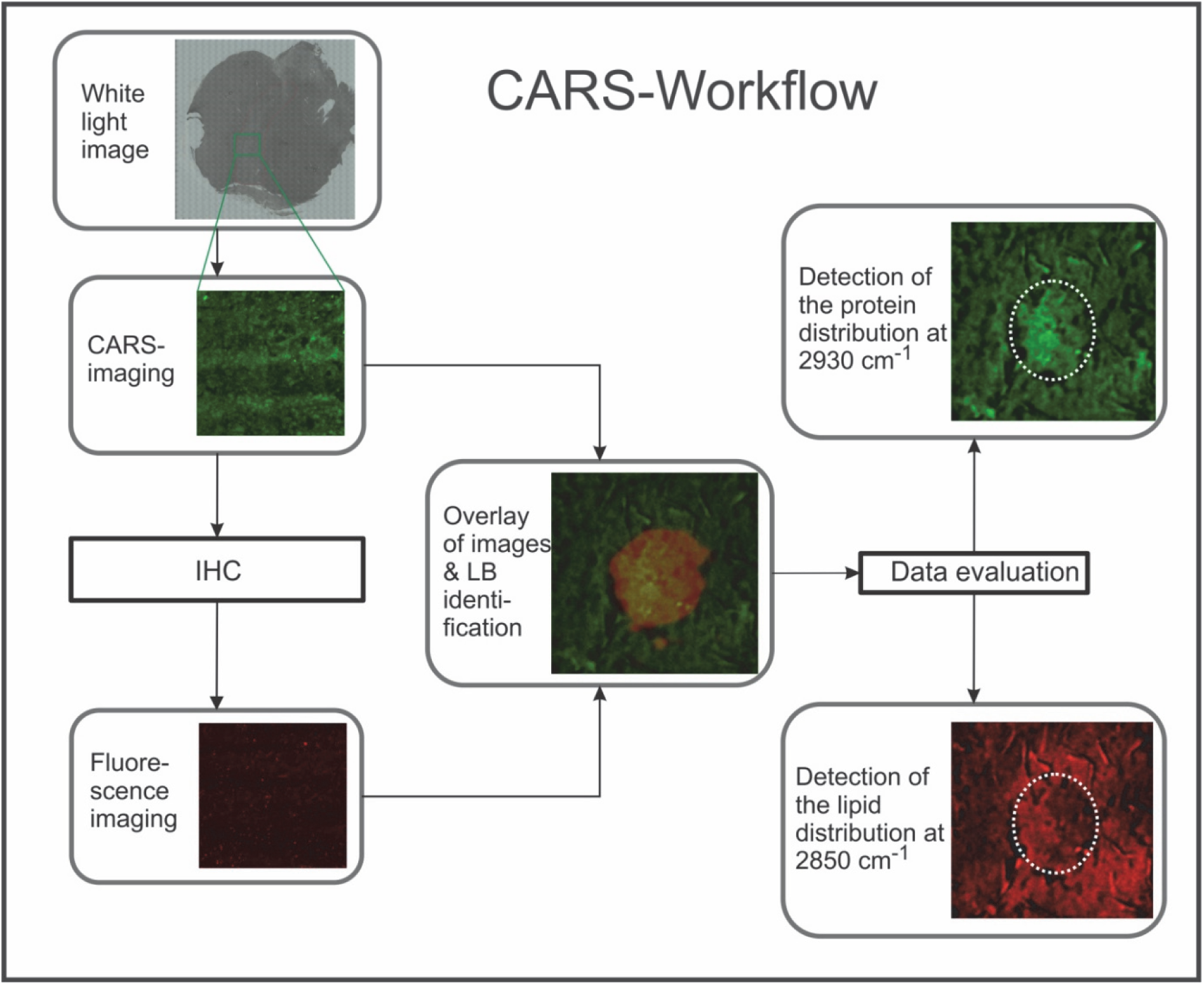
CARS-Workflow for the detection of the lipid and protein distribution in Lewy bodies. White light images were measured to detect the SN. The CARS-intensity was measured as overview images. On the same slide an immunohistological staining was performed and the fluorescence was detected for aSyn and Ser129-p aSyn. The images were manually overlaid. LBs were identified and small regions-of-interest (ROIs) were extracted. These ROIs were evaluated and the protein and lipid distribution of aSyn-positive inclusions was detected.

**Supplementary Figure 12:**
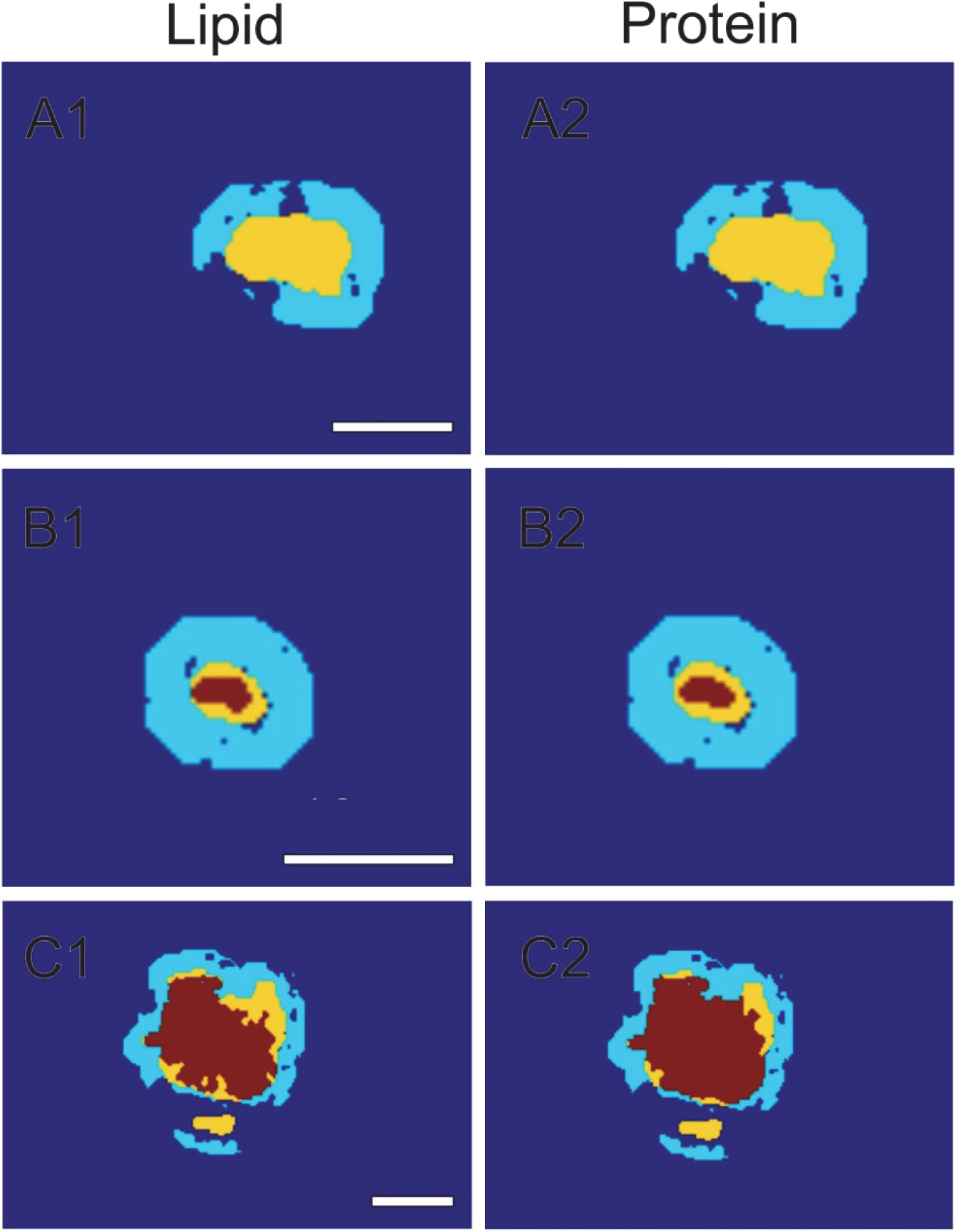
Evaluation of the centralization of lipids and proteins by CARS. The LBs were identified by aSyn staining (yellow) in the CARS-intensity images. For an objective evaluation of CARS intensity of the LBs, the mean CARS intensity of the direct surrounding (A, cyan), a donut with a width of 3.5 µm, was compared with the CARS-intensity of each LB pixel (A, yellow). CARS-pixel-intensities higher than 1.4 times the mean CARS-intensity of the surrounding were defined as higher protein/lipid content. The ratio between the CARS-pixel-intensities of the LB and the mean CARS intensity of the surrounding were calculated and areas with higher protein/lipid content were marked in red. Areas without CARS intensity (holes) were excluded by intensity thresholding in the first step. Scale bars = 10 µm.

## LegendSupplementary Video files

**Supplementary Video 1:** 3D reconstruction based on deconvolved CSLM images revealing the distribution of Ser129-p. 119CTT and 122CTT aSyn in an onion-skin type LB in the SN of patient PD7.

**Supplementary Video 2:** 3D reconstruction based on deconvolved CSLM images revealing the distribution of Ser129-p. 119CTT and 122CTT aSyn in another onion-skin type LB in the SN of another PD patient (PD6).

**Supplementary Video 3:** 3D reconstruction based on deconvolved CSLM images revealing the distribution of antibodies directed against NT, NAC domain and CT in a onion-skin type LB in the SN of patient PD8.

**Supplementary Video 4:** 3D reconstruction of an entire onion-skin type LB based on deconvolved CSLM images showing the distribution patterns of antibodies against different PTMs and domains of aSyn in the SN of patient PD1.

**Supplementary Video 5:** 3D reconstruction based on deconvolved CSLM images of an expansive-appearing aSyn inclusion showing the distribution patterns of antibodies against different PTMs and domains of aSyn in the SN of patient PD8.

**Supplementary Video 6:** 3D reconstruction based on deconvolved STED images showing the cage-like framework formed by Ser129-p aSyn and cytoskeletal components at the periphery of an onion-skin type LB in the SN of patient PD7.

**Supplementary Video 7:** z-stack visualization of a cytoskeletal framework at the periphery of nigral LBs in the SN of patient PD7 (deconvolved CSLM images)

**Supplementary Video 8:** z-stack visualization of a wheel-like structure of neurofilaments at the periphery of nigral LBs in the SN of patient PD6 (deconvolved CSLM images)

**Supplementary Video 9:** 3D reconstruction based on deconvolved CSLM images showing the Ser129-p aSyn immunopositive cytoplasmic network in an iLBD donor (iLBD1).

**Supplementary Video 10:** 3D reconstruction based on deconvolved STED images showing localization of a CTT-reactive punctae at the outer membrane of a mitochondrion immunopositive for VDAC/Porin in the hippocampus of patient PD4.

